# Turnover modulates the need for a cost of resistance in adaptive therapy

**DOI:** 10.1101/2020.01.22.914366

**Authors:** Maximilian Strobl, Jeffrey West, Yannick Viossat, Mehdi Damaghi, Mark Robertson-Tessi, Joel Brown, Robert Gatenby, Philip Maini, Alexander Anderson

**Affiliations:** Department of Integrated Mathematical Oncology, H. Lee Moffitt Cancer Center Tampa, USA; Wolfson Centre for Mathematical Biology, University of Oxford, Oxford, UK; Ceremade, Université Paris-Dauphine, PSL, Paris, France; Department of Cancer Physiology, H. Lee Moffitt Cancer Center Tampa, USA; Cancer Biology and Evolution Program, H. Lee Moffitt Cancer Center Tampa, USA

**Author notes:** Joint Senior Authors (Philip Maini, Alexander Anderson).

## Abstract

“Control and conquer” - this is the philosophy behind adaptive therapy, which seeks to exploit intra-tumoural competition to avoid, or at least, delay the emergence of therapy resistance in cancer. Motivated by promising results from theoretical, experimental and, most recently, a clinical study in prostate cancer, there is an increasing interest in extending this approach to other cancers. As such, it is urgent to understand the characteristics of a cancer which determine whether it will respond well to adaptive therapy, or not. A plausible candidate for such a selection criterion is the fitness cost of resistance. In this paper, we study a simple competition model between sensitive & resistant cell populations to investigate whether the presence of a cost is a necessary condition for adaptive therapy to extend the time to progression beyond that of a standard-of-care continuous therapy. We find that for tumours close to their environmental carrying capacity such a cost of resistance is not required. However, for tumours growing far from carrying capacity, a cost may be required to see meaningful gains. Notably, we show that in such cases it is important to consider the cell turnover in the tumour and we discuss its role in modulating the impact of a cost of resistance. Overall, our work helps to clarify under which circumstances adaptive therapy may be beneficial, and suggests that turnover may play an unexpectedly important role in the decision making process.

## Introduction

The evolution of drug resistance is one of the biggest challenges in cancer therapy. In 1947, Farber et al [1] for the first time reported that children suffering from acute leukaemia, a hitherto untreatable disease, showed “significant improvements” [1] upon treatment with a chemical agent, 4-amino-pteroylglutamic acid. However, they also also noted that these remissions were temporary [1]. This pattern still holds true today for many patients, and applies to both chemo-as well as targeted therapies.

Research aiming to combat drug-resistance has traditionally focussed on developing drugs which target either the resistance mechanism or kill the cell through a different route. As an alternative to this molecular approach, a number of authors (most notably Gatenby and colleagues [2, 3, 4, 5], but also [6], [7] and [8]) have proposed that resistance could be delayed, if not averted, through changes in drug-scheduling. The design principle behind current treatment schedules is to maximise cell kill by treating at the maximum tolerated dose (MTD) in as continuous a fashion as toxicity permits. However, if drug-resistant cells are present *a priori* or develop during treatment, such aggressive treatment releases these cells from the competition for space and resources and facilitates their growth, a process known as “competitive release” [2, 3, 7]. Inspired by approaches used in the management of invasive species and agricultural pests, Gatenby et al [2, 3] proposed adaptive therapy which aims not to eradicate the tumour, but to control it. Therapy is applied to reduce tumour burden to a tolerable level but is subsequently modulated or withdrawn to maintain a pool of drug-sensitive cancer cells. Over the past ten years, a number of studies have shown that adaptive therapy can extend time to progression (TTP) *in vivo*, in ovarian cancer [3], breast cancer [9], and most recently in melanoma [10]. Moreover, the interim analysis of the first human trial of adaptive therapy, applied to the treatment of metastatic, castrate-resistant prostate cancer with androgen-deprivation therapy, reported an increase in TTP of at least 10 months and a reduction in cumulative drug usage of 53% [11]. While it is challenging to prove that competition is the mechanism through which adaptive therapy extends TTP, and not, for example, synergistic effects on vasculature or the immune system, some evidence exists. Bacevic et al [12] showed that low dose treatment was able to minimise the frequency of resistant cells for longer than high dose treatment in a spheroid model of cyclin-dependent kinase inhibitor treatment in colorectal cancer, which has neither vasculature nor an immune system. Moreover, they found that this was not true if they repeated the experiment in a 2-D cell-culture model in which cells compete less strongly [12]. Furthermore, Smalley et al [10] observed that in both mouse and human samples of melanoma there was an enrichment for cells with a drug-sensitive transcriptional signature if the tumours had previously experienced a drug holiday, as would be predicted by the competitive control hypothesis.

The success in prostate cancer has spurred interest in extending adaptive therapy to other cancers such as thyroid cancer and melanoma (clinicaltrials.gov identifiers NCT03630120 and NCT03543969, respectively). As such, it is urgent to understand the characteristics of a cancer that determine whether it will respond well to adaptive therapy, or not. Bacevic et al [12] showed through *in vitro* experiments and an agent-based computational model that it is important that cells are spatially constrained, as otherwise the competition is too weak to effectively control the growth of resistant cells. Furthermore, through ODE modelling they found that the fitness of the resistant population when the population is rare is a key determinant of the benefit derived from adaptive therapy [12]. These results were corroborated by Gallaher et al [13] who compared two adaptive therapy strategies on an agent-based model which modelled resistance as a continuous trait. They found that the initial abundance of resistance, the rate of spatial mixing through migration, and the rate of acquisition of resistance through mutation, were the key factors in determining the benefit from adaptive therapy, and which adaptive therapy strategy was most effective. Using a non-spatial game theory model, West et al [14] showed that from a mathematical perspective the benefit of adaptive therapy is determined by the relative position of the initial tumour to the zero-growth isocline of the resistant population in the state space of cell type frequencies. Biologically, this means that the stronger the selection against resistance in the absence of drug, the longer adaptive therapy can be expected to extend TTP. Finally, while not directly focussed on cancer, Hansen et al [8] present a general argument for how to decide when treatment should aim to control and when it should aim to eradicate a pathogen. They derive quantitative expressions for the levels of initial resistance above or below which a management strategy such as adaptive therapy would perform better than an eradication strategy [8].

However, a factor which is still poorly understood is the role of the cost of resistance. *“Cost of resistance”* means that a resistance adaptation confers a fitness advantage under drug exposure, but it comes at a fitness cost in a treatment-free environment [2, 15]. It is a phenomenon which has been widely studied in agricultural pests and antibiotic resistance [16, 17] and is typically experimentally measured in one of three ways: i) Mono-culture experiments, which measure the growth rate of the resistant population in the absence of drug in isolation, ii) Competition experiments, which study the relative frequency of sensitive and resistant cells over time when cultured together, and iii) Reversal experiments, which explore how long it takes for resistant cells to lose their resistance mechanism if grown in the absence of drug (Figure 1A-C). For example, Gallaher et al [13] report that the doubling time of doxorubicin-resistant MCF7Dox breast cancer cells is decreased by 50% compared to their sensitive counterparts, and that in competition experiments the sensitive cells outcompete the resistant cells in the absence of the drug. This is likely because these cells use P-Glycoprotein-related efflux pumps to resist treatment, for which they have to divert energy away from proliferation towards running of the pumps [13, 18]. Bacevic et al make similar observations for CDKi resistant cells and a number of further examples can be found in the literature (e.g. [19, 20, 21, 22, 23]). In their original work on adaptive therapy, Gatenby et al [2, 3] used the cost of resistance to motivate adaptive therapy, and most theoretical studies since have made this assumption [12, 13, 14]. However, not all resistance mechanisms come at a cost. Behrens et al [24] find that A2780 cisplatin-resistant ovarian cancer cells have a shorter doubling time than the sensitive parental line (22.1h compared to 25.3h). Similar observations have been made for certain colorectal cancer cell lines [21], and Kaznatcheev et al [23] report that even though their resistant cells grow more slowly in mono-culture, their growth is supported by the presence of sensitive cells in co-culture [23]. In addition, the impact of a cost is context dependent. When we repeat the experiments by Gallaher et al [13] in 3-D spheroids, we find that drug-resistant MCF7 cells now grow faster than their sensitive counterparts (Figure 1D). However, when we reduce the glucose concentration in the culture medium, this advantage is lost, a finding also made by Silva et al [4] in 2-D culture (Figure 1E). Given this wide range of possibilities for how a resistant population might, or might not, differ from their sensitive counterparts, it is important to clarify the relationship between cost of resistance and the success of adaptive therapy.

**Figure 1:**
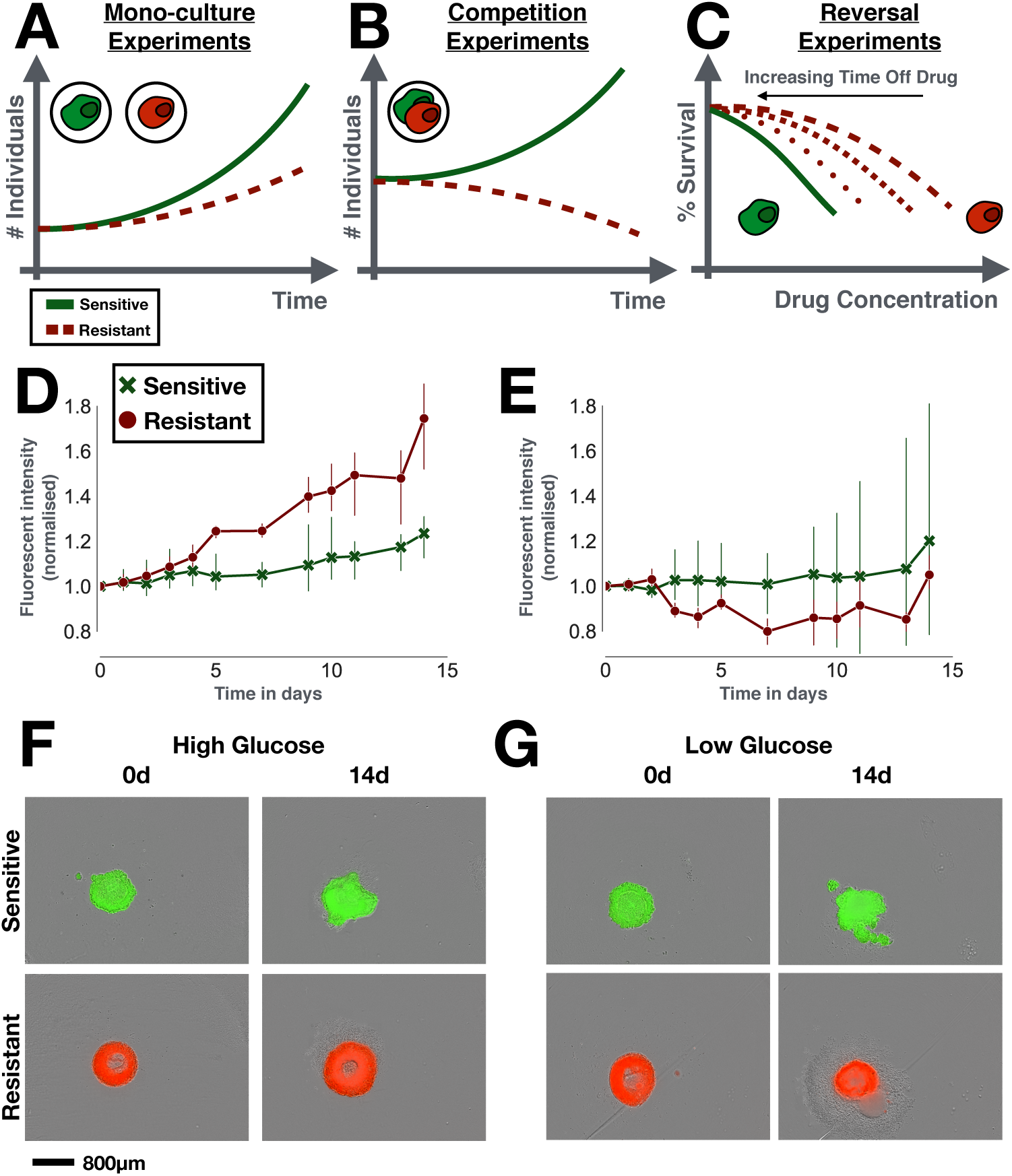
Resistance costs in theory and in practice. A-C) The three main experimental designs used to test for a cost of resistance (all done in drug-free conditions). A) Mono-culture experiments test for changes in growth rate, size, migration rate etc of the resistant strain in isolation. B) Competition experiments compare the abundance of sensitive and resistant cells in co-culture over time. C) Reversal experiments examine the rate at which drug resistance is lost, if the drug is withdrawn. To do so, the resistant population is cultured in a drug-free environment, and its drug response is tested at regular time intervals. D-G) In vitro spheroid experiments comparing the growth of doxorubicin-sensitive and resistant MCF7 cells in mono-culture. Sensitive and resistant cells were GFP and RFP tagged, respectively. D) In normal medium, resistant cells grow faster than sensitive cells, showing that resistance does not necessarily have to be costly (3 replicates per group). E) Under glucose starvation this advantage is lost (3 replicates per group). This shows that the environmental context has to be considered when studying fitness costs. F & G) Example images of the initial (0d) and final time points (14d) from D & E showing the fluorescent signal overlayed on the bright-field image. For experimental details see Section S1 in the Supplementary Data.

The aim of this paper is to investigate the impact of a cost of resistance on adaptive therapy. We use a simple mathematical model in which we divide the tumour into drug-sensitive and drug-resistant cells and model their growth with two ordinary differential equations (ODEs). We compare the TTP under standard-of-care continuous therapy with that of the adaptive therapy algorithm used in the clinical trial by Zhang et al [11], first in the absence, and subsequently in the presence, of a cost of resistance. We will show how increased tumour density and small levels of pre-existing resistance maximise competition between sensitive and resistant cell populations within the tumour, making adaptive therapy superior to continuous therapy. Subsequently, we will demonstrate that high cell turnover is a key factor to consider, not only to understand the impact of a cost of resistance, but also to assess the ability to control a tumour with adaptive therapy more generally.

## Methods

### Mathematical model

Tumours are heterogeneous populations of cells with differential responses to drug, indicating a degree of pre-existing resistance in most tumours [25, 26, 27]. To model this heterogeneity, we assume two competing cell types: drug-sensitive cells, *S*(*t*), and fully resistant cells, *R*(*t*), modelled via the following equations:

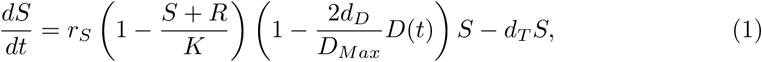

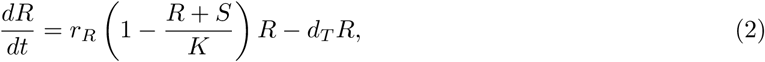

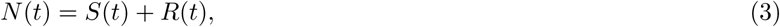

with initial conditions:

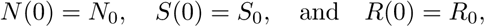

where *N*_0_ = *S*_0_ + *R*_0_. The model is derived based on the following assumptions:

- In isolation each population grows logistically with proliferation rates *r*_*S*_ and *r*_*R*_, where the fraction of dividing cells decays linearly from 1 to 0 as the population approaches its environmental carrying capacity, *K*. Furthermore, cells die at a density-independent rate *d*_*T*_, where for simplicity we assume that this turnover rate is the same for both populations.
- Cells compete for resources and space according to the Lotka-Volterra competition model. This means that the presence of the competitor reduces a population’s growth rate in a fashion that is linearly proportional to the competitor’s population density. For simplicity we will assume that inter- and intra-species competition coefficients are identical, and equal to one.
- Only actively dividing cells are killed by the drug. Many chemotherapies induce DNA damage or inhibit the cell division machinery, which induces apoptosis only in cells that attempt to divide [28]. We will adopt the classical Norton-Simon model [29] which assumes that cell kill increases proportionally with the fraction of dividing cells and linearly with the drug dose, *D*(*t*), so that at MTD a fraction *d*_*D*_ of dividing cells are killed.
- To facilitate our analysis, we make the simplifying assumption that the cost of resistance manifests itself solely in the growth rate *r*_*R*_. This is also the most common way in which the cost is modelled (e.g. [12, 13, 26]).

Note that the factor of 2 in the drug response accounts for the fact that if a cell dies during mitosis not only the potential daughter but also the mother are lost. The curious reader may refer to the Supplementary Data for a mathematical discussion of the steady states of this system, (see Section S4) and a discussion of the implications of other manifestations of a cost of resistance (see Section S6).

We will consider two treatment schedules:

1. Continuous therapy at the MTD, *D*_*Max*_: *D*(*t*) = *D*_*Max*_∀ *t*.
2. Adaptive therapy which withdraws treatment once a 50% decrease from the initial tumour size is achieved, and reinstates it once the original tumour size (*N*_0_) is reached:

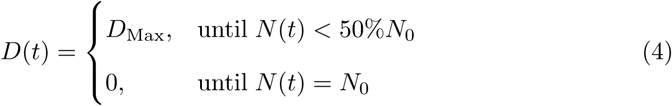

In order to facilitate numerical simulation of the model, we non-dimensionalise Equations (1) - (3) using *r*_*S*_ as a time scale, and *K* as a scale for the cell densities (Table 1). For details of the non-dimensionalisation and the numerical methods, see Section S2 of the Supplementary Data.

**Table 1:** Mathematical model parameters and their ranges.

| Parameter | Description | Value | Reference |
| --- | --- | --- | --- |
| $s = S/K$ | Sensitive cell density (normalised) | 0.0 - 1.0 | |
| $r = R/K$ | Resistant cell density (normalised) | 0.0 - 1.0 | |
| $r_S$ | Sensitive cell proliferation rate and model time scale | $0.027d^{-1}$ | Adopted from [11]. |
| $\hat{r}_R := \frac{r_R}{r_S}$ | Resistant cell proliferation rate (normalised) | 0.5 - 1.0 | Lower limit: [13];<br>Upper limit: assumption of no cost |
| $\hat{d}_T := \frac{d_T}{r_S}$ | Cell turnover rate (normalised) | 0.0 - 0.5 | Lower limit: assumption of no turnover;<br>Upper limit: see Section S3. |
| $\hat{d}_D := 2d_D$ | Drug-induced cell kill factor (see text for further explanation) | 1.5 | Adopted from [30]. |
| $n_0 := \frac{N_0}{K}$ | Initial tumour cell density (normalised) | 0.1 - 0.75 | Parameter sweep; values within this range reported by [31]. |
| $f_R := \frac{R_0}{N_0}$ | Initial resistant cell fraction | 0.001 - 0.1 | Parameter sweep; values within this range reported by [27]. |
| $f_S := \frac{S_0}{N_0}$ | Initial sensitive cell fraction | 0.9 - 0.999 | Determined by $1 - f_R$. |

Note that the factor of 2 in the drug response accounts for the fact that if a cell dies during mitosis not only the potential daughter but also the mother are lost. The curious reader may refer to the Supplementary Data for a mathematical discussion of the steady states of this system, (see Section S4) and a discussion of the implications of other manifestations of a cost of resistance (see Section S6).

### Parameterising the model

Given the key role which prostate cancer has played in the development of adaptive therapy we parametrise our model according to this disease. As such, we adopt the proliferation rate for sensitive cells given in [11] (*r*_*S*_ = 0.027d^−1^) as our time scale and the drug kill parameter 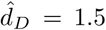 from [30]. For the other parameters we perform parameter sweeps within their biologically realistic ranges. All parameters, their definitions and their ranges are summarised in Table 1.

## Results

In advanced cancers, curative approaches rarely show durable complete response. Instead, treatment success is often defined by how long therapy can prevent the tumour from progressing beyond a certain size (i.e. TTP). Herein we compare the model-predicted TTP for both adaptive (TTP_*AT*_) and continuous therapy (TTP_*CT*_), respectively, using RECIST criteria (20% increase in tumour size from the pre-treatment baseline). We show that time gained by adaptive therapy depends on the following tumour characteristics: initial tumour density (*n*_0_; see Figure 3), initial levels of pre-existing resistance (*f*_*R*_; see Figure 3), cost of resistance (*r*_*R*_ < *r*_*S*_; see Figure 4), and the density-independent cell turnover rate (*d*_*T*_; see Figure 5).

**Figure 2:**
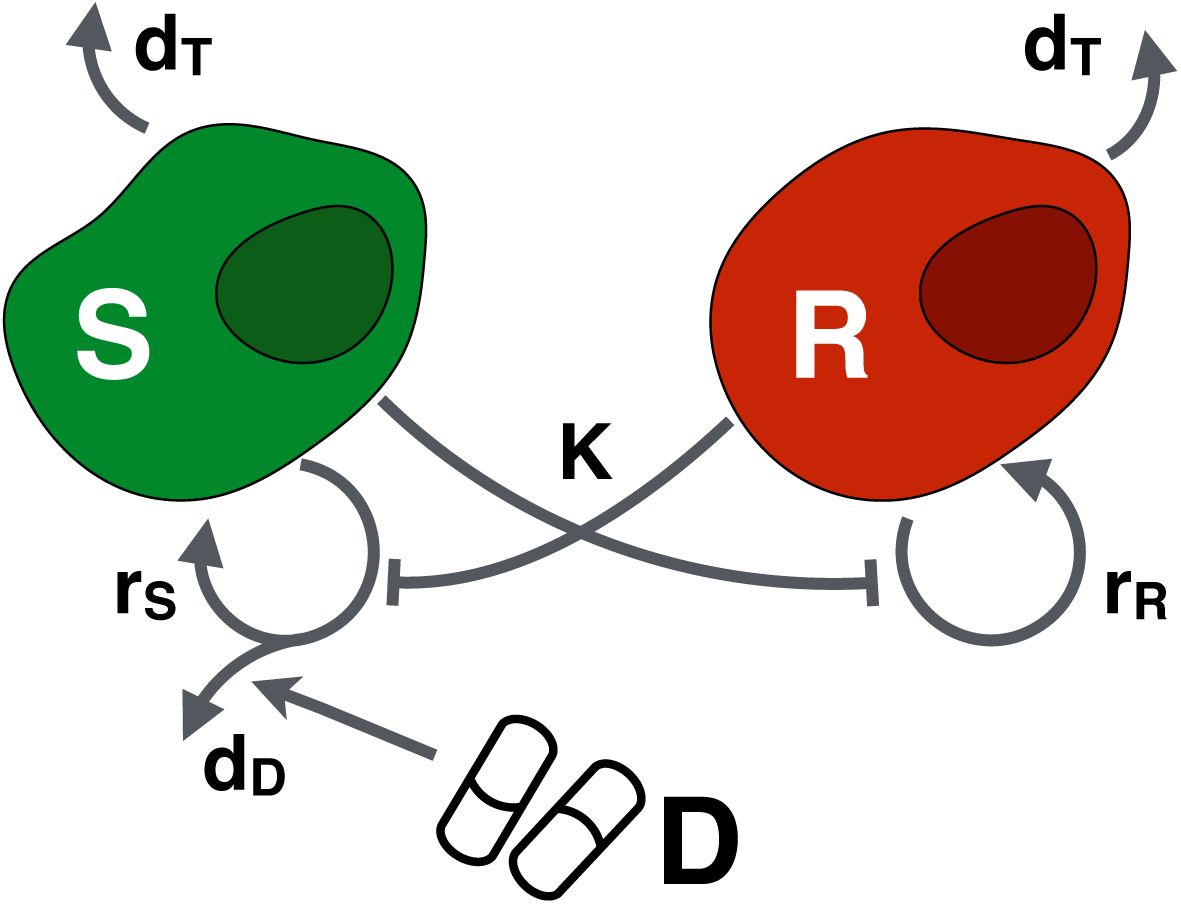
The mathematical model. Drug-sensitive (S) and resistant cells (R) proliferate at rates, rS and rR, respectively, and die at rate dT. Proliferating sensitive cells die at a rate dD when exposed to drug, D. Finally, both populations compete for resources, where K denotes the total (shared) environmental carrying capacity.

**Figure 3:**
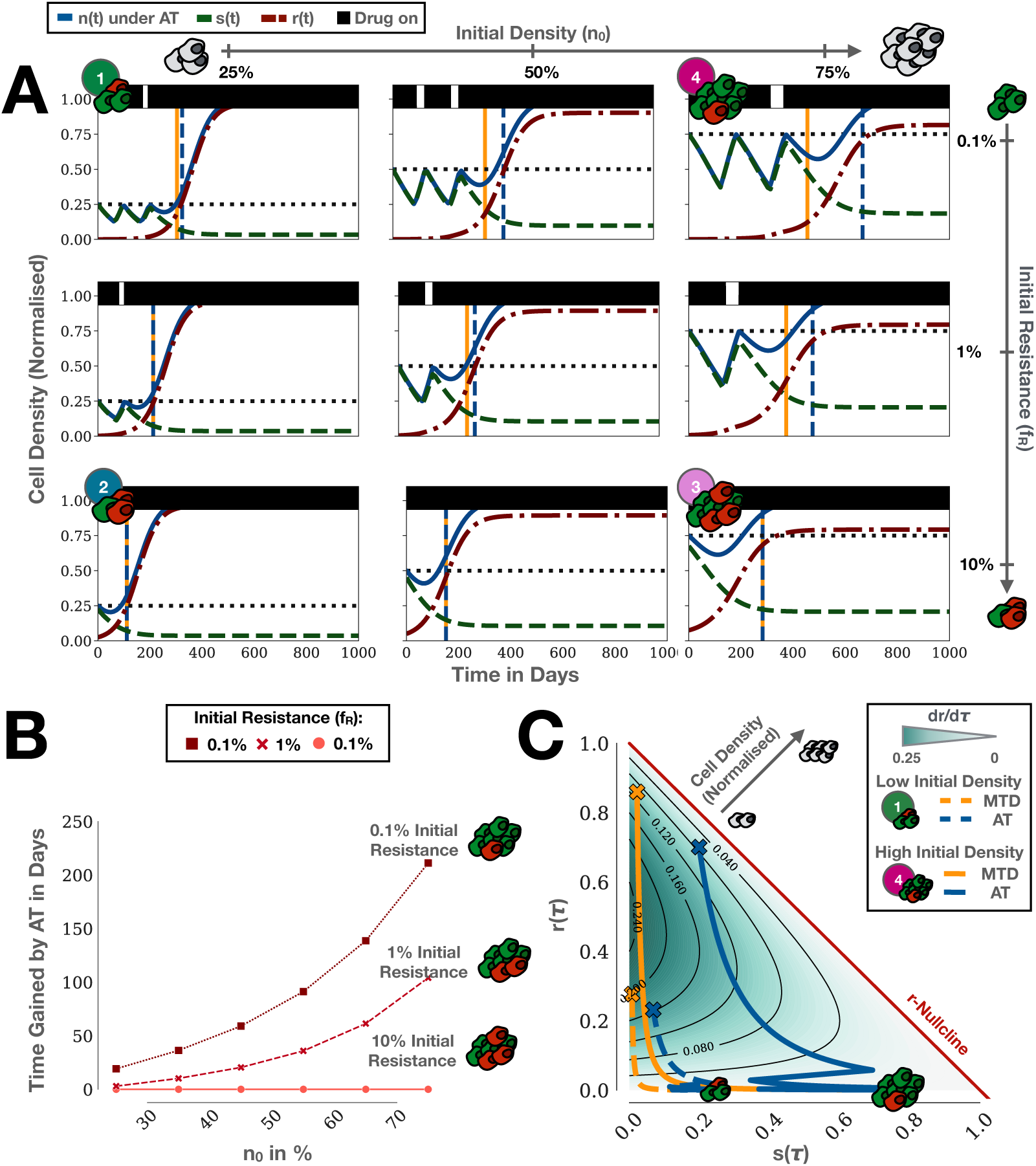
Adaptive therapy (AT) can extend TTP compared to continuous therapy even in the absence of a cost of resistance. For simplicity, we are assuming no turnover here 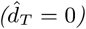. A) Simulations of adaptive therapy for a cohort of tumours with different initial compositions. Vertical dashed lines indicate the time of progression of continuous (yellow) and adaptive therapy (blue), respectively. B) Gain in TTP by adaptive compared to continuous therapy as a function of initial proximity to K and abundance of resistance. C) dr/dτ as a function of s and r, together with treatment trajectories for Tumours 1 ((n_0_, f_R_) = (25%, 0.1%)) & 4 ((n_0_, f_R_) = (75%, 0.1%)) from A. Crosses indicate progression. This shows how adaptive therapy extends TTP by minimising dr/dτ via competition, and demonstrates that for certain tumours adaptive therapy can extend TTP even if no cost of resistance is present.

**Figure 4:**
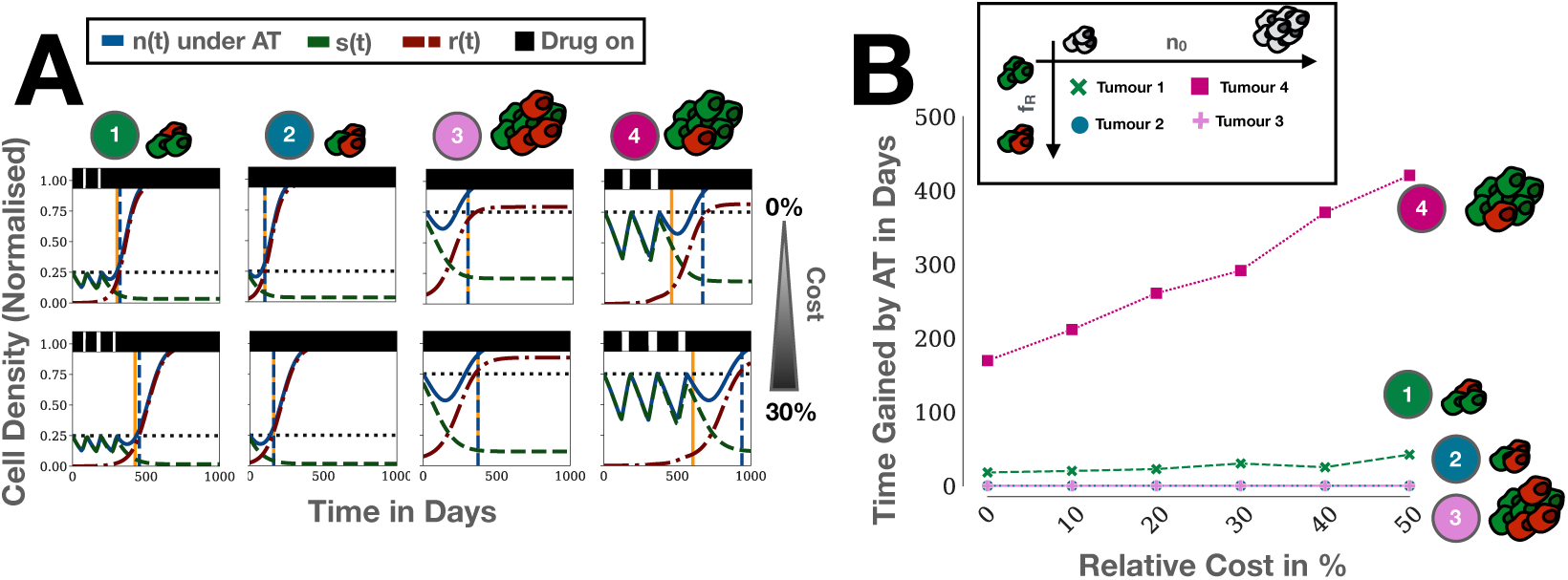
In the absence of turnover 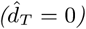 a cost of resistance enhances adaptive therapy only when resistance is rare and tumours are close to carrying capacity. A) Simulations of Tumours 1-4 from Figure 3A with and without a 30% cost of resistance (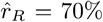; Tumours 1 & 4 given in Figure 3; Tumour 2: (n_0_, f_R_) = (25%, 10%); Tumour 3: (n_0_, f_R_) = (25%, 10%)). B) Gain in TTP by adaptive therapy compared to continuous therapy for different levels of cost of resistance for Tumours 1-4.

**Figure 5:**
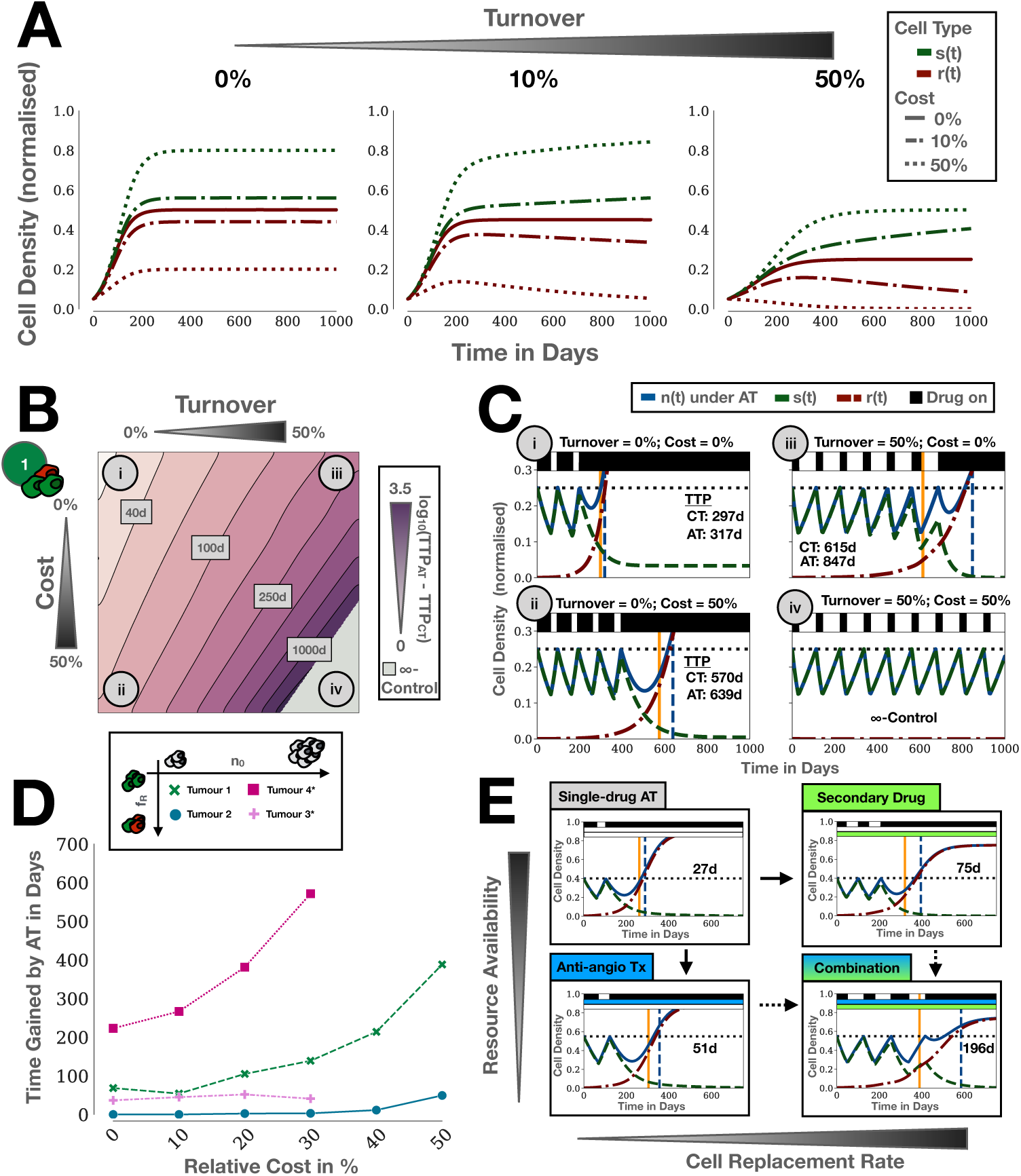
Turnover as a driver of adaptive therapy. A: Simulation of a competition experiment (no drug; n_0_ = 10%, f_R_ = 50%). In the absence of turnover, even a significant cost does not result in the elimination of the resistant cells. In contrast, in its presence, cells paying a resistance cost are eventually outcompeted by sensitive cells. B: TTP gained by adaptive therapy as a function of cost and turnover for Tumour 1. Turnover increases the benefit of adaptive therapy and amplifies the effect of a cost. C: Simulations of the treatment dynamics for four combinations of cost and turnover corresponding to the four case studies highlighted in B. D: Turnover increases TTP gained by adaptive therapy (TTP_AT_-TTP_CT_) both in the presence and absence of a resistance cost for a range of different tumour compositions (Tumours 1 and 2 as in Figure 3A; Tumour 3*: (n_0_, f_R_) = (50%, 10%); Tumour 4*: (n_0_, f_R_) = (50%, 0.1%); above a cost of 30% Tumours 3* and 4* become indefinitely controllable and so no TTP can be obtained). E: Improving tumour control by increasing competition with a second drug which either reduces K (e.g. anti-angiogenic drug) or increases d_T_ /r_R_ (e.g. low dose chemotherapy; for corresponding values of the intensity of competition see Figure S4). Inset numbers give TTP gained by adaptive therapy. Single drug adaptive therapy: n_0_ = 40%, f_R_ = 1%, 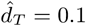; Anti-angiogenic Tx changes n_0_ to 55%; Secondary drug increases 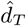 to 0.25.

**Figure 6:**
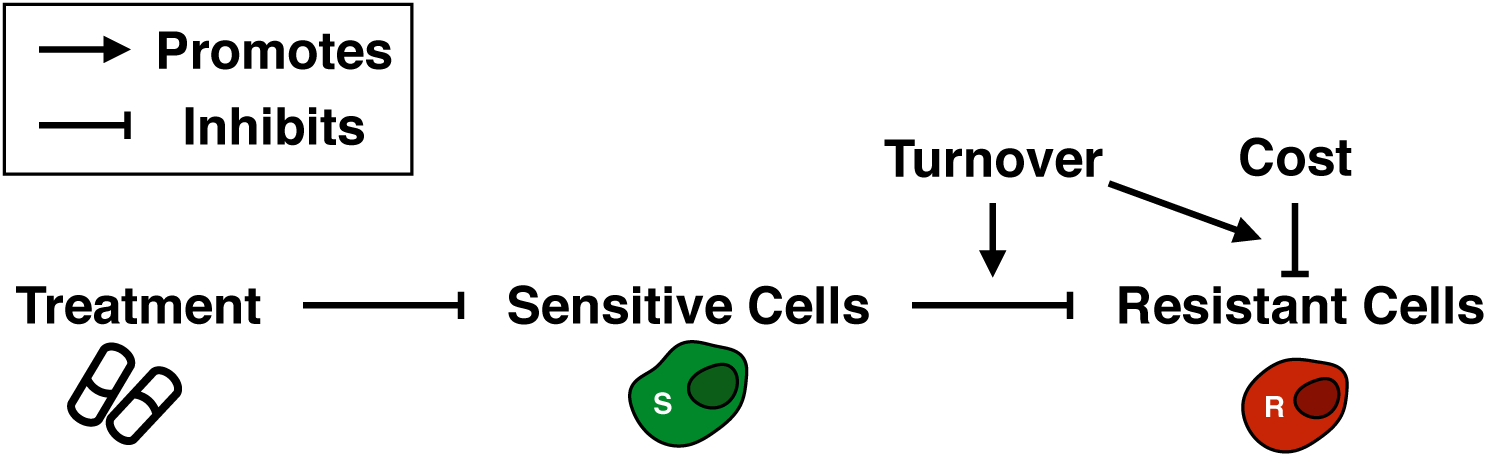
The interplay between cost of resistance and turnover in adaptive therapy. Sensitive cells slow the growth of resistant cells through competition, but are killed by treatment. A cost of resistance reduces the resistant growth rate. Turnover amplifies the impact of both a cost of resistance and the strength competition.

### Adaptive therapy extends TTP even without a resistance cost

In the absence of a resistance cost, adaptive therapy still results in significant improvements in TTP over continuous therapy. Figure 3A shows how these gains are modulated by tumour density (plots, left to right), and pre-existing resistance (top to bottom). Each subpanel shows sensitive (red lines), resistant (green) and total (blue) populations over time under adaptive therapy dosing. Treatment administration is illustrated by black bars at the top of the graphs. Figure 3A clearly shows how the TTP of adaptive therapy (vertical blue dashed lines) outlasts the TTP of continuous therapy (vertical yellow) for the combination of low resistance and high tumour density (top right panel). Figure 3B quantifies time gained (TTP_*AT*_ - TTP_*CT*_) for a range of tumour density and resistance values. For our parameter sweep, we find that adaptive therapy can extend TTP by between 3 and 104 days when the initial resistance fraction is *f*_*R*_ = 1%, and by between 19 and 211 days when *f*_*R*_ = 0.1%. We also note that in the worst case scenario (*f*_*R*_ = 10%), adaptive therapy becomes indistinguishable from continuous therapy, so that while there is no benefit, the patient has also not progressed faster than under standard-of-care (bottom row in Figure 3A and Figure 3B).

### Adaptive therapy treatment vacations provide a benefit only if intra-tumoural competition is strong

Moreover, we find that each of the two characteristics of the tumour’s initial composition has a distinct impact on the treatment dynamics. As we increase the initial abundance of resistant cells from 0.1% to 10%, we decrease the number of completed adaptive therapy cycles (Figure 3A). In the most extreme case, at 10% initial resistance, treatment can not decrease the tumour burden sufficiently to trigger any treatment withdrawal in the three tumours. In contrast, increasing the initial tumour density, which biologically corresponds to greater competition of resources, does not alter the adaptive therapy cycle number, but does increase the benefit delivered by each cycle. For example, even though all tumours with 1% initial resistance complete one adaptive therapy cycle, a meaningful benefit in TTP is only achieved in the case when the tumour is 75% saturated (Figure 3A). The reason for this is that the competition exerted by sensitive cells only has significant impact on the growth of the resistant cells if the tumour is close to *K*. In summary, for adaptive therapy to provide a benefit the tumour burden must undergo a sufficient decline to allow for treatment withdrawal (small *f*_*R*_), and competition within the tumour must be sufficiently strong to noticeably slow the expansion of the resistant population (proximity to *K*).

### Adaptive therapy extends TTP by minimising the resistant population growth rate

How can we explain the benefit of adaptive over continuous therapy in the absence of a cost of resistance? We apply the framework by Hansen et al [8] to derive the following explanation. Progression is primarily driven by the expansion of the resistant population, as drug-sensitive cells are easily depleted by additional treatment. Thus, the more a treatment strategy can inhibit the resistant population growth rate, *dR/dt*, whilst also maintaining control of the tumour size overall, the longer TTP. To illustrate this, we plot *dr/dτ* (the non-dimensional form of *dR/dt*) as a function of the tumour composition in Figure 3C. In this representation a tumour lesion is seen as a point in a two dimensional space, where its x-position represents the current relative density of sensitive cells, *s*(*τ*), and its y-position the current density of resistant cells, *r*(*τ*). Each point is coloured according to the resistant population growth rate. This representation clearly illustrates how high tumour densities (see inset grey arrow) are generally associated with lower resistant growth rates. Re-writing Equation (2) as:

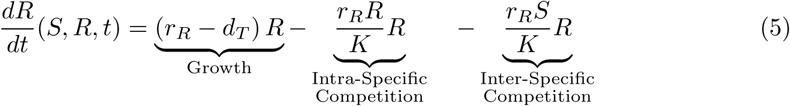

shows that this slow-down occurs due to intra- and inter-specific competition. Importantly, while a cost in *r*_*R*_ modulates the strength of growth inhibition through competition, it is not required (as also illustrated in Figure 3C).

During therapy, the composition of a tumour changes, so that treatment corresponds to a trajectory through the *S*−*R* space. In Figure 3C we show trajectories of continuous (yellow) and adaptive therapy (blue) for two tumours corresponding to Tumours 1 & 4 in Figure 3A. As can be seen, continuous therapy trajectories tend to traverse regions of high resistant growth (dark green shading). In contrast, these same regions are avoided under adaptive therapy regimens, especially for Tumour 4. Again we can formalise this using Equation (2): If *S*_*CT*_ (*t*) and *S*_*AT*_ (*t*) denote the density of sensitive cells after *t* subsequent days of treatment under continuous and adaptive therapy, respectively, then we have that *S*_*CT*_ (*t*) ≤ *S*_*AT*_ (*t*) because continuous therapy does not provide sensitive cells with an opportunity to recover. Assuming that the tumour is still far from progression (*R* ≪ *S*) so that the resistant cells primarily compete with sensitive cells, the resistant growth rate is greater for continuous 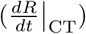 than for adaptive therapy 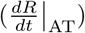:

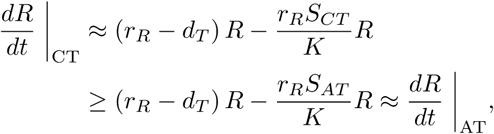

This implies that adaptive therapy can provide a benefit even in the absence of a cost of resistance. This benefit increases with increased tumour density as *S*_*AT*_ */K* can be maintained large for longer. In addition, this implies that the most effective adaptive therapy would maintain the tumour at its original size for as long as possible in order to maximise the effects of inter-specific competition.

### In the absence of turnover, cost of resistance increases time gained by adaptive therapy only when resistant cells are initially rare

Next, we examine the role of a resistance cost. Figure 4 compares low density tumours (Tumour 1 and 2, repeated from Figure 3) with high density tumours (Tumour 3 and 4). As expected, TTP_AT_ (blue line) increases with a cost of resistance in all four cases (Figure 4A). However, an increase of the benefit of adaptive over continuous therapy is only seen when tumours are close to *K* and resistant cells are rare, as in Tumours 1 and 4. In contrast, in highly resistant tumours, a cost of resistance increases the TTP in roughly equal terms for continuous and adaptive therapy. This effect is consistent for a wide range of resistance cost values (Figure 4B).

### Turnover mediates the impact of a cost of resistance

So far we have assumed that the turnover of tumour cells is negligible. However, tumours are subject to resource starvation and immune predation, resulting in continuous tumour turnover. In Figure 5A we show how such turnover modulates the impact of a cost of resistance on the competition between sensitive and resistant cells in the absence of drug. Increasing the cost of resistance (dashed lines) leads to lower levels of resistance in untreated populations. Importantly, while a cost of resistance reduces the number of resistant cells, these cells never go extinct (Figure 5A). In contrast, if we introduce intrinsic cellular turnover 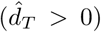, Figure 5A shows that resistant cells go extinct over time. We conclude that selection against resistance depends not only on the cost of resistance but also the turnover rate.

How does this insight affect adaptive therapy? In Figure 5B we show TTP_AT_ as a function of turnover and cost for Tumour 1. If turnover is low, a large cost of resistance results only in small gains for adaptive therapy, as seen by comparing Cases i and ii in Figures 5B & C. In contrast, if turnover is high, adaptive therapy provides significant benefits even if the resistance cost is small, or completely absent (Case i vs iii in Figures 5B & C). We also observe in Case iv that once turnover increases beyond a threshold of 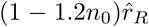 the tumour becomes indefinitely controllable in the model, so that continuous and adaptive therapy maintain the tumour below the size for progression forever (see Section S5 for further discussion).

Next, we examine how this result generalises as we change the initial tumour density and resistance fraction. We repeat the cost-gain relationship analysis from Figure 4B with a turnover rate of 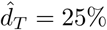. As Tumours 3 & 4 are indefinitely controllable in this parameter regime (see Figure S3C), we replace them by tumours with a slightly lower initial density (*n*_0_ = 50%), denoted by Tumours 3* & 4*, respectively. We find that all but the less dense and highly resistant tumour (Tumour 2) benefit from adaptive therapy even in the absence of a resistance cost (Figure 5D). Interestingly, Tumour 3* did not benefit from adaptive therapy in the absence of turnover (Figure 3A; bottom row, centre panel), but with turnover, adaptive therapy becomes superior to continuous therapy. Moreover, in all cases a cost of resistance now increases the time gained with adaptive therapy. This shows that turnover increases the gains of adaptive therapy and amplifies the effect of a cost of resistance, thereby relaxing the need for the tumour to be close to carrying capacity for effective adaptive therapy. The initial resistance fraction, however, remains an important determinant of TTP_AT_.

### Resource availability, turnover and cost combine to determine competition in the tumour

Why does turnover increase the benefit of adaptive therapy? In Figure 3A we showed that the closer the tumour is to its carrying capacity at the start of treatment, the greater the benefit of adaptive therapy. This is because a larger value of *N*_0_*/K* corresponds to greater resource limitation and competition. To investigate the impact of turnover on competition, we rewrite Equation (2) as:

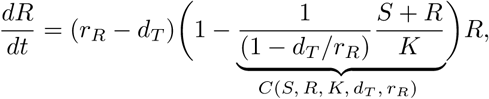

where *C*(*S, R, K, d*_*T*_, *r*_*R*_) describes the growth inhibition of the resistant population through intra- and inter-specific competition. A value of *C* = 0 corresponds to no inhibition, whereas a value of *C* = 1 results in the complete growth arrest of the resistant population. We see that the strength of competition is determined by three factors: i) The available resources, *K*; ii) the density of cells, *S* + *R*; and iii) the ratio of turnover to proliferation, *d*_*T*_ */r*_*R*_, known as the cell replacement rate. Thus, an increased cell replacement rate - due to either increased turnover or a proliferative cost - results in stronger competition (see also Figure S4). Alternatively, one can say that the resistant population’s growth is limited not by the environmental carrying capacity, *K*, but by the effective carrying capacity *K*_Eff_ = (1 − *d*_*T*_ */r*_*R*_)*K*.

### Improving adaptive therapy by amplifying competition

Based on our results we hypothesise that one may improve the efficacy of adaptive therapy through administration of additional treatments to increase cellular competition in one of two ways. Firstly, one may reduce the environmental carrying capacity, *K*, for example, by administering an anti-angiogenic drug. Secondly, one may increase the cell replacement rate, *d*_*T*_ */r*_*R*_. For example, if the primary drug which is given adaptively is a targeted therapy, this secondary drug could be a low dose chemotherapy. In Figure 5E we illustrate how these additional therapies can increase competition and consequently significantly extend tumour control under adaptive therapy.

## Discussion

With four clinical trials of adaptive therapy already ongoing (clinicaltrials.gov identifiers: NCT02415621; NCT03511196; NCT03543969; NCT03630120) and more in preparation, it is important to develop criteria to identify which patients might benefit from adaptive therapy over standard MTD approaches. Intuitively, it appears that the presence of a fitness cost of resistance would be a great facilitator of adaptive therapy, and should be considered as an inclusion criterion. However, as Bacevic et al [12] and Gallaher et al [13] show, the presence of a cost does not guarantee that the resistant population can be effectively controlled. Moreover, the experimental work which motivated the trial in prostate cancer did not find a cost of resistance *in vitro*, yet adaptive therapy has extended median TTP by over 10 months [11].

The aim of this paper is to consolidate these findings and develop an understanding of the circumstances under which the presence of a cost of resistance is required for adaptive therapy to provide a benefit, and under which circumstances it is not. To do so, we developed a simple two-population Lotka-Volterra competition model and treated it according to the adaptive therapy algorithm from [11]. We have shown that the time gained compared to continuous therapy depends on three key aspects: i) the initial fraction of resistant cells (*f*_*R*_), ii) the proximity of the tumour to environmental carrying capacity (*n*_0_), and iii) the rate of cellular turnover (*d*_*T*_). Previously it has been demonstrated that the smaller the fraction of resistance in a tumour and the closer it is to carrying capacity, the greater the benefit of adaptive relative to continuous therapy [8, 12, 13]. We extend these findings by showing that for such tumours a cost of resistance is not necessary for adaptive therapy to be beneficial. While its presence increases the time gained by adaptive therapy, our work suggests that significant benefits may be obtained even in its absence. Some of our results can be proven rigorously and in a broader context, at least in the absence of turnover. These details are beyond the scope of this paper, but can be found in [32].

Since adaptive therapy aims to exploit competition between cells it makes sense that its efficacy increases the closer the tumour is to carrying capacity. However, what is the meaning of carrying capacity in a dynamic environment such as a tumour? Commonly, carrying capacity, *K*, is interpreted as the maximum population size the environment can support due to resource constraints and is thought of as independent of the population’s intrinsic growth rate, *r* (see, for example, these three recent cancer publications in the *Bulletin of Mathematical Biology*: [33, 34, 35]). However, biological populations have intrinsic turnover, and therefore the population’s actual equilibrium will be smaller than *K*. Moreover, this equilibrium is dependent on *r*, so that a cost in *r* changes not only the exponential growth rate of the population but also its equilibrium value - the shorter the life span of a cell, the fewer opportunities it will have to divide, and the greater will be the impact of interference through intra- and inter-specific competition. This fact resolves a number of commonly criticised issues of the logistic growth law and helps to translate results from ecological to evolutionary theory [36, 37, 38].

These observations regarding turnover have implications for the success of adaptive therapy (Figure 5A). When cell death is negligible, a cost of resistance at best slows the expansion of resistance, but can not prevent it. In contrast, if there is significant turnover, even a small cost of resistance may make the resistant population controllable for a long time. In essence, turnover modulates the intensity of selection against resistance during the drug-free intervals. As such, careful thought needs to be given to choose appropriate experimental model systems to test adaptive therapy. For example, Bjorkman et al [39] have shown in bacteria that different fitness-compensating mutations are selected depending on whether the bacteria are evolved *in vitro* or *in vivo*. In cancer, Bacevic et al [12] found that resistance could be contained through competition in a spheroid, but not in a 2-D cell culture model. While *in vitro* no resistance cost may be observed, tumour control may still be possible *in vivo* or in patients as turnover, via greater nutrient deprivation, and immune suppression will intensify competition, and amplify even small fitness differences.

What is the turnover rate of tumour cells? While this is challenging to quantify in patients, the existing evidence paints a rather dynamic picture of growing tumours. Many cancers exhibit areas of necrosis, a sign that conditions within a tumour often cause cell death in certain areas. Furthermore, the doubling time expected from the fraction of dividing cells seen in histological analyses, and the actually observed volumetric doubling time show substantial discrepancies, suggesting the rate of cell death in tumours closely matches that of cell production (see Section S3 for further discussion; [40, 41, 42, 43]). It is also important to remember that in homeostatic tissue, birth and death are in fact balanced, as the tissue continuously renews itself without changing in size [42]. We propose that the speed of turnover in a tumour should be further quantified and, in combination with the homeostatic turnover rate in its tissue of origin, should be investigated as selection criteria for adaptive therapy.

While we focussed our attention on the impact of a cost in the proliferation rate, it is clear that it may also manifest itself in other ways. For example, in the 3-D spheroids we consider in Figure 1, there is significantly more debris around the resistant spheroids than around their sensitive counterparts (see also S1.4). Also, the necrotic core appears enhanced (Figure 1F & G). This suggests that the cost in this case might manifest itself in an increased turnover rate. We also show that our analysis easily extends to costs in the turnover rate and carrying capacity, where either costs in growth rate or costs in the carrying capacity will have the greatest impact depending on the cell replacement rate (Section S6).

We make a number of simplifications in our model. Firstly, similar to [8, 12] we assume that the tumour is not curable. However, as can be seen in Figure 5D, including a cost of resistance and greater turnover also result in fewer cells at the nadir during continuous therapy. This implies that the tumours in which adaptive therapy will be most effective are those in which continuous therapy just misses being curative, an observation similar to that made in [7] and [13]. Thus, the very long gains predicted under adaptive therapy have to be viewed with some scepticism, since in a proportion of these patients continuous therapy would have cured the tumour. That being said, adaptive therapy is intended for advanced settings in which the available treatment is rarely curative, which suggests that such cases will be rare [2, 5]. Moreover, it seems plausible that if one finds that adaptive therapy controls the tumour very well, one could then decide to switch to a curative approach. In fact, adaptive therapy has been shown to have stabilising effects on tumour vasculature [9], which suggests that a curative approach may be even more effective after an initial period of adaptive therapy.

Furthermore, we do not consider the potential health-risks associated with the increased tumour burden under adaptive therapy. As a result, our model predicts that the longest TTP can be achieved by keeping the tumour as close to its initial size as possible. This is true also for most previous studies carried out in this area ([3, 10, 12, 13]). As Hansen et al [8] and Viossat and Noble [32] show, if one accounts for potential *de novo* resistance acquisition, this high burden can generate situations in which maintaining a pool of sensitive cells can in fact accelerate the emergence of the resistant population. Investigating the risks associated with the higher average tumour burden under adaptive therapy is an important area of future research.

Finally, in order to facilitate analysis we neglect the impact of space in this manuscript. However, both [12] and [13] have previously demonstrated that in the absence of turnover the spatial architecture of the tumour is a key determinant of the success of adaptive therapy. If resistant cells can be “trapped” by sensitive cells in the centre of the tumour, the tumour may be controlled for a long time [12, 13]. Going forward it would be important to assess whether our conclusions extend to a spatial setting. While turnover will increase the selection against resistant cells, it might also facilitate their escape by creating gaps in the layer of sensitive cells surrounding them. Interestingly, using an extension of their model in [13], Gallaher et al [44] have already shown that turnover mediates the proliferation-migration trade-off in a tumour. In addition, while we define *N*(*t*) to represent tumour cell density, we treat it effectively as a volume in determining progression. As a consequence, we have to assume that the tumours we consider are below a certain density, so that we can still observe progression. This both restricts us to lower tumour densities than what might be observed in actual patients and can result in the prediction of indefinite TTP in certain parameter regimes. Also this issue could be addressed more accurately with a spatial model.

To conclude, we want to highlight interesting parallels and possible opportunities from the pest and antibiotic resistance management literature, which have investigated fitness costs of resistance since the mid-20^th^ century (see, for example, [45] for an early review of cost of resistance in insecticides). Firstly, we note that similar to cancer it has always been a controversial issue. For example, Bergelson et al [46], find that out of 88 studies examining the cost question in plants, 44 show a cost, 40 find no difference and 4 even observe a benefit to resistance. Similar patterns can be seen for pesticide resistance [45, 47] and antibiotic resistance [16, 17]. Whether a cost is present depends on the resistance mechanism, and its environmental as well as genetic context [15, 17, 46]. Thus, in developing adaptive therapy we should not assume that a cost of resistance is always present or is uniform throughout the tumour, or that it will remain constant over time. That being said, in line with the predictions of this paper, resistance management can be successful even if the specific variations of the cost of resistance are uncertain or absent. For example, the resistance management scheme for the insecticide producing *Bacillus thuringensis* (Bt) crop has been successful despite inconclusive evidence regarding the presence of a resistance cost [47, 48, 49].

In order to compensate for the fact that there is usually more than one evolutionary trajectory to resistance, many pest resistance management strategies include multiple treatment modalities [50, 51]. As such, we advocate multi-drug approaches in which one, or several drugs, are given adaptively. Initial theoretical work on multi-drug adaptive therapy has already been carried out [5, 30, 52], and we have illustrated here how by targeting the resource availability or turnover in a tumour with secondary drugs one can greatly enhance tumour control. We also note that while abiraterone was given adaptively in the initial prostate cancer adaptive therapy trial (NCT02415621), patients were concurrently receiving a continuous dose of Lupron which suppresses systemic testosterone production and reduces the cancer’s supply of growth factors. A follow-up study now gives both Abiraterone and Lupron in an adaptive fashion (NCT03511196). Going forward it will be important to extend this work to develop strategies which exploit a cost of resistance, yet are robust if this cost disappears due to environmental or genetic compensation. With better understanding of tumour growth, resistance costs, and turnover rates, adaptive therapy can be more carefully tailored to patients who stand to benefit from it the most.

## Acknowledgements

We acknowledge Dr. Carlo Maley for inspiring Figure 6 with a sketch he presented at the Meeting on Evolutionary Therapy on the 24th of May 2018 (https://twitter.com/evoltherapy/status/999665282737205249?s=21). This research was supported by funding from the Engineering and Physical Sciences Research Council (EPSRC) and the Medical Research Council (MRC) [grant number EP/L016044/1]. Viossat benefited from the support of the FMJH “Program Gaspard Monge for optimization and operation research and their interactions with data science” and from the support from EDF, Thales, Orange and Criteo. Anderson & Gatenby gratefully acknowledge funding from both the Cancer Systems Biology Consortium and the Physical Sciences Oncology Network at the National Cancer Institute, through grants U01CA232382 and U54CA193489 as well as support from the Moffitt Center of Excellence for Evolutionary Therapy.

