## supplementary data for "Turnover modulates the need for a cost of resistance in adaptive therapy"

March 17, 2020

#### S1 *In vitro* spheroid experiments

##### S1.1 Sensitive and resistant cell culture

MCF7 cells were acquired from American Type Culture Collection (ATCC, Manassas, VA, 2007–2010) and were maintained in RPMI 1640 (Life Technologies) supplemented with 10% FBS (HyClone Laboratories). Cells were tested for mycoplasma contamination and authenticated using short tandem repeat (STR) DNA typing according to ATCC guidelines. To establish stable cell lines expressing GFP and RFP, the MCF-7 cells were infected with Plasmids expressing RFP or GFP using Fugene 6 (Promega) at an early passage and were selected using 2 $\mu$g/ml puromycin (Sigma). During this experiment MCF7-GFP cells were kept always sensitive and MCF7-RFP resistant to Doxorubicin. To make the resistant cells we grew the MCF7-RFP in 0.1 mM Doxorubicin for three months. MCF7-RFP-DOX cells keep their resistant phenotype after freeze thaw or for several passages in regular media (not shown).

##### S1.2 3D spheroid co-culture

Perfecta3® 96-well Hanging Drop Plates and non-adhesive U-Shape Bottom 96-well plates were used to grow the primary spheres containing 20,000 cells total. Two experiments were conducted: 1) MCF7-GFP and MCF7-RFP-DOX cells were grown in mono-culture in high glucose conditions (10mM), and 2) both cells lines were grown in mono-culture in low glucose conditions (1mM). In each case the experiment had a starting density of 20,000 cells and physiological (7.4) pH. Each experiment had three replicates. An Incucyte microscope placed in 37 degree and 5% $CO_2$ was used to image the spheroid growth every 24h over 14 days.

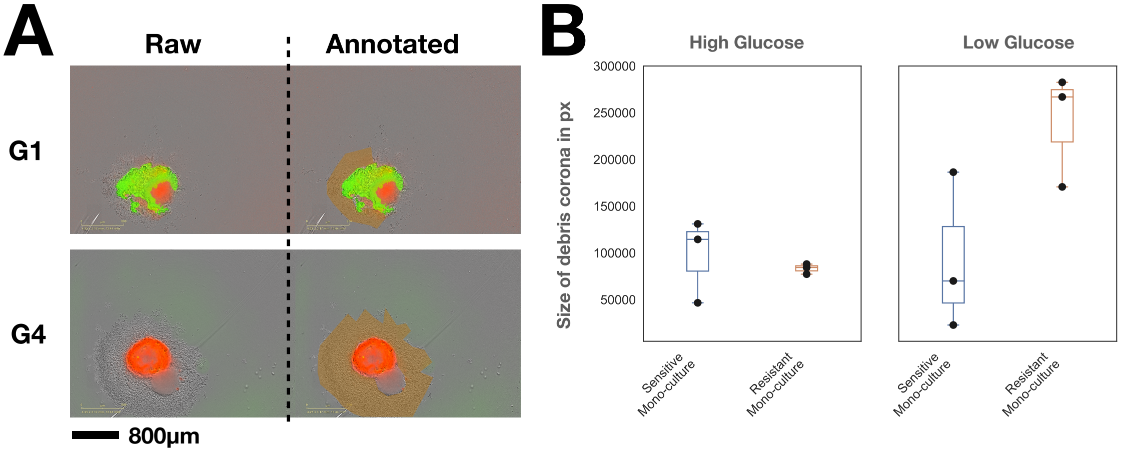


Figure S1: *Doxorubicin-resistant MCF7 cells (MCF7-RFP-DOX) show increased sensitivity to glucose starvation at 14d compared to their drug-sensitive counterpart (MCF7-GFP). A) Microscopy images with overlayed fluorescence illustrating the segmented debris corona for two wells (G1: sensitive mono-culture in low glucose conditions; G4: resistant mono-culture in low glucose conditions). In addition, this exemplifies the background noise (red signal in G1, green signal in G4) which has been removed from the data in Figure 1G. B) Comparison of the size of the debris corona between sensitive and resistant spheroids. While there is no difference in high glucose conditions (two-sample t-test:* $t_{3,3} = 0.54$, $p_{S>R} = 0.3$*), the difference is significant in low glucose conditions (two-sample t-test:* $t_{3,3} = 2.45$, $p_{S>R} = 0.035$*).*

#### S1.3 Image Analysis

Images and fluorescent intensity were extracted using in the Incucyte built-in software. Relative fluorescent units (RFUs) were normalized at each time point by averaging red/green fluorescence when the corresponding cell type was absent then subtracting this value from other fluorescent values of that cell type. To assess relative growth over time, RFUs were normalised relative to the fluorescence recorded 6h after seeding of the spheroids (6h so to allow for some settling-in time of the culture). All analysis and post-processing were carried out in Python 3.6.

#### S1.4 Analysis of Cell Turnover

In order to quantify the build-up of debris around the spheroids over time we manually segmented the corona around each spheroid at the final time point. The size of the corona in pixel is shown in Figure S1. As can be seen, the corona is noticeably larger around MCF7-RFP-DOX spheroids than around MCF7-GFP spheroids in low glucose conditions (Figure S1A).

### S2 Non-dimensionalisation & Numerical Methods

#### S2.1 Non-Dimensionalisation

In order to reduce the number of free variables, we non-dimensionalise Equations (1) - (3). Given that we are interested in studying the treatment dynamics if we change the characteristics of the resistant cells relative to the sensitive cells, we base our scales on the sensitive population. Specifically, we will use the following transformations:

$$\tau = r_S t, \quad s = \frac{S}{K}, \quad r = \frac{R}{K}, \quad c = \frac{D}{D_{\text{Max}}}, \quad \text{and} \quad n = \frac{N}{K}.$$

This yields:

$$\frac{ds}{d\tau} = (1 - s - r)\,(1 - \hat{d}_D\, c(\tau))\, s - \hat{d}_T s, \tag{1}$$

$$\frac{dr}{d\tau} = \hat{r}_R\,(1 - s - r)\, r - \hat{d}_T r, \tag{2}$$

$$n(\tau) = s(\tau) + r(\tau), \tag{3}$$

where:

$$\hat{d}_D = 2d_D, \quad \hat{d}_T = \frac{d_T}{r_S}, \quad \text{and} \quad \hat{r}_R = \frac{r_R}{r_S}.$$

The initial conditions transform to:

$$n(0) = \frac{N_0}{K} := n_0, \quad s(0) = \frac{S_0}{K} := s_0, \quad \text{and} \quad r(0) = \frac{R_0}{K} := r_0,$$

and the treatment schedules are adjusted accordingly as well (not shown). Finally, for notational convenience we will define $f_S := S_0/N_0$ and $f_R := R_0/N_0$ as the initial fractions of sensitive and resistant cells, respectively.

#### S2.2 Numerical Methods

The equations were solved using the RK5(4) explicit Runge-Kutta scheme [1] provided in Scipy (specifically, the SCIPY.INTEGRATE.ODE class). Adaptive therapy was implemented by simulating an interval $[t, t + \Delta t]$ at a time, and subsequently updating the dose for the next interval according to the algorithm in Equation 4. In preliminary experiments we tested different values of $\Delta t$ and found $\Delta t = 1$d to provide a good trade-off between speed and accuracy (not shown). All simulations were carried out in Python 3.6, using Scipy 1.1.0 and Numpy 1.15.1. Visualisations were produced with Pandas 0.23.4, Matplotlib 2.2.3, and Seaborn 0.9.0. The code will be made available on-line upon publication.

| Pathological Type | $T_\text{V}$ (in days) (95% CI) | $T_\text{Pot}$ (in days) | Estimated Growth Fraction (in %) | Cell Loss Factor (in %) |
|---|---|---|---|---|
| Embryonal tumours | 27 (22-33) | 1.66 | 90 | 94 |
| Hemato-sarcoma | 29 (23-37) | 1.7 | 90 | 94 |
| Mesenchymal Sarcomas | 41 (35-50) | 13.2 | 11 | 68 |
| Squamous cell carcinoma | 58 (48-70) | 6.0 | 25 | 90 |
| Adeno-carcinoma | 83 (72-96) | 23.8 | 6 | 71 |

Table A1: *Proliferation rates and cell loss factors measured in human tumours (taken from the meta-analysis by Malaise and colleagues in [2]).*

#### S3 Justification of the choice of turnover rates

While cell turnover due to apoptosis, necrosis and immune predation is recognised as a feature of many tumours, little is known about the rate at which tumour cells die. Most available data infer the amount of cell loss indirectly using a method introduced by Gordon Steel in the 1960s [3, 4, 5]. The idea is to estimate the theoretically possible ("potential") doubling time, $T_\text{Pot}$, achieved in the absence of any cell death, and the actually observed volumetric doubling time of a tumour, $T_\text{V}$. By comparing these values one can compute the "cell loss factor", $\phi$, defined as $\phi = 1 - T_\text{Pot}/T_\text{V}$ which represents the relative rate of cell death to proliferation (assuming an exponentially growing tumour). The greater the value of $\phi$, the greater the loss of cells due to cell death, predation, or emigration [3, 4, 5].

$T_\text{V}$ is estimated from x-ray, CT or MRI images. $T_\text{Pot}$ is obtained by measuring the fraction of cells in S-phase in the tumour via halogenated pyrimidine, bromo- or iododeoxyuridine labelling. Cells carrying out DNA synthesis incorporate these agents into their DNA, and can subsequently be identified using histological techniques or flow cytometry. The proportion of labelled cells is termed the labelling index (LI). To obtain $T_\text{Pot}$, the LI is adjusted to account for proliferating cells which are not currently in S-phase and is multiplied by an estimate of the cell cycle time, based on the time required for DNA synthesis ($T_S$). This gives $T_\text{Pot} = \lambda T_S/LI$, where $\lambda$ denotes the proportion of the cell cycle spent in the S-phase. For a detailed review of the process of estimating $T_\text{Pot}$ see [6].

In Table A1 we show estimates for the cell loss factor for a range of human tumours. These are all well in the excess of 50% - an observation which has been corroborated by a large number of studies over the years (see [2, 4, 7] for extensive reviews). To translate estimates for $\phi$ to values of our model parameter $\hat{d}_T$ we need to note that $\phi$ compares the cell turnover rate to the effective proliferation rate of the population (which takes into account the actual fraction of proliferating cells), not the intrinsic growth rate. To illustrate this, consider a logistic model of a growing tumour. Let $N(t)$ denote the total number of cells and assume that cells divide and die at rates $r_T$ and $d_T$, respectively, so that: $dN/dt = r_T(1 - N/K)N - d_T N$. If we were to measure this tumour at time $t^*$ we would observe a growth fraction of $(1 - N(t^*)/K)$ and estimate (making the assumption of the Steel method that this growth fraction remains constant), that $T_{\text{Pot}} = \ln(2)/r_T(1 - N(t^*)/K)$. The actual volumetric doubling time (again assuming the growth fraction remains constant) would be $T_{\text{V}} = \ln(2)/(r_T(1 - N(t^*)/K) - d_T)$. Thus, $\phi = d_T/(r_T(1 - N(t^*)/K))$. To obtain $\hat{d}_T := d_T/r_T$, we therefore want to scale the observed values of $\phi$ by the fraction of dividing cells. Applying this calculation to the data in Table A1 we obtain values between 4% and 81%, with rates for solid tumours (the settings in which adaptive therapy has been most studied in) clustered below 25%. In light of this uncertainty, we choose to consider values of up to 50%. If turnover rates are higher this would simply further exaggerate our results. Future research should investigate turnover rates in tumours in more detail to obtain more accurate estimates.

### S4 Steady State Analysis

For completeness we here analyse the more general model discussed in Section S6 of which the model which forms the focus of the main paper is a special case. Also, so to explicitly show the relationship of the steady states with the system parameters we will here work with the dimensional form of the equations. We will begin with a general discussion before providing two specific examples illustrating the key features of the phase plane dynamics which are driving our results. Assuming continuous therapy at dose $D(t) = D^*$ Equations (5)-(6) have the following steady states:

- Tumour Elimination (SS1): $(S^*, R^*) = (0, 0)$. A linear stability analysis gives eigenvalues $\lambda_1 = r_S(1 - d_D D^*) - d_S$ and $\lambda_2 = r_R - d_R > 0$, which shows that this state is always unstable and so, unless there are no resistant cells in the tumour or $d_R > r_R$, tumour elimination is not possible in this model. This is because there is no mechanism in this model to prevent the outgrowth of resistant cells except for inhibition from sensitive cells.

- A Fully Sensitive Tumour (SS2): $(S^*, R^*) = \left(\left(1 - \frac{d_S}{r_S(1-d_D D^*)}\right) K_S, 0\right)$ which corresponds to a tumour consisting entirely of sensitive cells. This state has eigenvalues $\lambda_1 = r_R - d_R - r_R \left(1 - \frac{d_S}{r_S(1-d_D D^*)}\right) \frac{K_S}{K_R}$ and $\lambda_2 = r_S(d_D D^* - 1)\left(1 - \frac{d_S}{r_S(1-d_D D^*)}\right)$, so that its feasibility and

| State | Description | A) No Turnover ($d_T = 0$) | B) With Turnover ($0 < d_T < r_S, r_R$) |
|---|---|---|---|
| SS1 | *Tumour Elimination* | $\boldsymbol{\lambda_1 = r_S(1 - d_D D^*)}$<br>$\lambda_2 = r_R$ | $\boldsymbol{\lambda_1 = r_S(1 - d_D D^*) - d_T}$<br>$\lambda_2 = r_R - d_T$ |
| SS2 | *Sensitive Tumour* | $\lambda_1 = 0$<br>$\boldsymbol{\lambda_2 = r_S(d_D D^* - 1)}$ | $\boldsymbol{\lambda_1 = d_T \left(\frac{r_R}{r_S(1-d_D D^*)} - 1\right)}$<br>$\boldsymbol{\lambda_2 = r_S(d_D D^* - 1)\left(1 - \frac{d_T}{r_S(1-d_D D^*)}\right)}$ |
| SS3 | *Resistant Tumour* | $\lambda_1 = 0$<br>$\boldsymbol{\lambda_2 = -r_R}$ | $\boldsymbol{\lambda_1 = d_T \left(\frac{r_S}{r_R}(1 - d_D D^*) - 1\right)}$<br>$\boldsymbol{\lambda_2 = d_T - r_R}$ |
| SS4 | *Coexistence* | No simple form available | No simple form available |

Table A2: *Eigenvalues for the cases considered in Figure S2 (assuming $d_D D^* \neq 1$). **Bolded values** indicate eigenvalues which are negative in at least part of the parameter space.*

stability depend on the effective drug kill rate $d_D D^*$, and the ability of the resistant cells to invade the equilibrium, which is determined by the balance of terms expressed in $\lambda_1$.

- A Fully Resistant Tumour (SS3): $(S^*, R^*) = \left(0, \left(1 - \frac{d_R}{r_R}\right) K_R\right)$ which describes a tumour consisting entirely of drug resistant cells. This state has eigenvalues $\lambda_1 = r_S(1 - d_D D^*)\left(1 + \left(\frac{d_R}{r_R} - 1\right)\frac{K_R}{K_S}\right) - d_S$ and $\lambda_2 = d_R - r_R < 0$, which implies its stability is dependent on the sign of $\lambda_1$.

- Coexistence (SS4): $(S^*, R^*) = \left(S, \left(1 - \frac{d_R}{r_R}\right) K_R - S\right)$, which describes a set of tumours consisting of a mixture of sensitive and resistant cells. This set only exists if the two non-zero nullclines precisely overlap (see Figure S2A for an illustration). No simple analytical form is available for the corresponding eigenvalues, and thus, they will be omitted here.

Having discussed the eigenvalues in general we will briefly show four specific examples to illustrate the impact of the turnover and a cost of resistance on the phase plane dynamics. To do so, we will again adopt the simplifying assumptions that $d_S = d_R := d_T$ and $K_S = K_R := K$. Unless otherwise stated simulations were done with the parameters from Table 1.

#### S4.1 No turnover

Assuming $d_T = 0$, we obtain the eigenvalues shown in Table A2. We note that $r_R$ has no effect on the position of the steady states and merely scales the eigenvalues so that a cost of resistance will have no influence on the structure of the phase space in this case. Thus, to simplify the following analysis we will assume there is no cost ($r_R = r_S$).

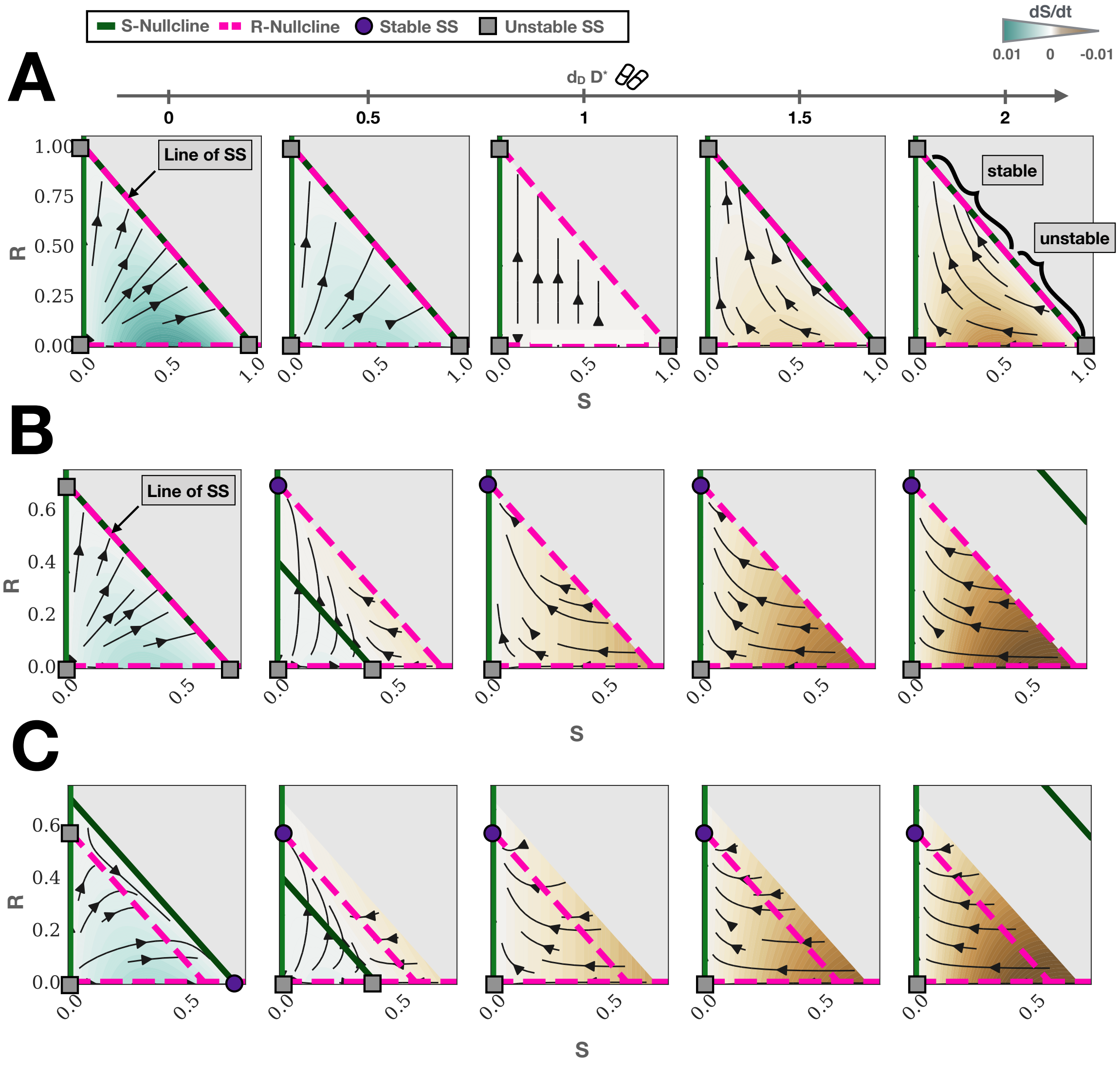


Figure S2: *Phase planes of the model (Equations (1) and (2)) under continuous therapy with increasing levels of drug kill, $d_D D^*$ (parameters as in Table 1). We assume no cost of resistance ($r_R = r_S$). SS stands for steady state. Grey area corresponds to tumours which are excluded from the analysis because they would shrink even without treatment. A) No turnover and no cost ($d_T = 0$, $r_R = r_S$). There is a continuum of steady states (SS4) given by $S + R = 1$. For $d_D D^* < 1$ the tumour continues to grow until it reaches carrying capacity on SS4. For $d_D D^* > 1$ sections of SS4 become unstable and we see the formation of heteroclinic orbits between unstable and the stable section of SS4. This represents tumours which are largely sensitive so that they will initially experience tumour shrinkage, but will subsequently recur due to the outgrowth of resistance. B) Turnover but no cost ($d_T = 0.1$, $r_R = r_S$). Unless $d_D D^* = 0$, the tumour becomes fully resistant over time. C) Turnover and cost ($d_T = 0.3$, $r_R = 0.7$). Provided the tumour is treated at low dose ($d_D D^* < 1 - r_R/r_S$), it can be maintained fully sensitive and decreased in size (not shown). If the dose is escalated beyond this threshold, the tumour will evolve to become fully resistant.*

Both SS2 and SS3 have zero eigenvalues because they lie on the manifold of steady states given by SS4. By considering the phase flow of Equations (1) & (2) one can see that between $d_D D^* = 0$ and $d_D D^* = 1$, the tumour will keep growing and converge to a point on SS4 (Figure S2A). The greater the value of $d_D D^*$, the larger the proportion of resistant cells at equilibrium. When $d_D D^* = 1$, $dS/dt$ becomes exactly 0 everywhere (Figure S2A; see also Equation 1). As $d_D D^*$ is increased above 1, the sign of $dS/dt$ switches (Figure S2A). Moreover, parts of the line SS4 become unstable, and we observe the formation of heteroclinic orbits connecting the unstable portions of SS4 to the stable portions of SS4 (Figure S2A). Biologically, the transition through $d_D D^* = 1$ represents a transition from when the drug only reduces the net growth of the population to when it actually causes decreases in the population size. The heteroclinic orbits represent tumours which initially shrink in size but subsequently recur. Interestingly, the fully resistant state becomes the globally absorbing state only as $d_D D^* \to \infty$. This is because the resistant cells protect the sensitive cell by saturating the environment, thereby preventing them to divide which makes them no longer sensitive to the drug.

#### S4.2 With turnover

If we assume that turnover is present ($0 < d_T < r_S, r_R$) we obtain the eigenvalues given in the right-hand side column of Table 1. To begin with, let us assume that there is no cost of resistance ($r_R = r_S$). If $d_D D^* = 0$, then we again have a line of steady states, but this time the tumour saturates below the carrying capacity, $K$ (Figure S2B). Importantly, if the drug kill is increased, then the S-nullcline given in dimensional form by $S(t) = \left(1 - \frac{d_T}{(1-d_D D^*)r_S}\right) K - R$, is now translated downwards. As a result, for $0 < d_D D^* < 1$, SS4 disappears and SS3 becomes stable ($\lambda_1, \lambda_2 < 0$) whereas SS2 is unstable ($\lambda_1 > 0$; see also Figure S2B). Thus, in contrast to when turnover is absent, in the presence of turnover the tumour evolves to become fully resistant even if treated at a very low dose (Figure S2B). When $d_D D^*$ is increased above 1, SS2 takes on a value above K which converges to $K$ from above as $d_D D^* \to \infty$ (see the expression for SS2 in Section S4 and Figure S2B). Whilst positive, this state is not biologically realistic as it implies that the population can grow beyond the environmental carrying capacity.

Finally, we consider the impact of a cost of resistance in this setting ($r_R < r_S$). From Table A2 we can see that provided $d_D D^* < 1 - r_S/r_R$, SS2 is stable whereas SS3 is unstable. Thus, in the presence of a cost low dose treatment can shrink the tumour whilst maintaining it stably sensitive (Figure S2C). However, if the drug kill is intensified beyond $1 - r_S/r_R$ SS2 becomes unstable and SS3 becomes stable (Table A2) and the tumour will become fully resistant over time (Figure S2C).

#### S5 Indefinite Tumour Control

In order for a tumour to progress in our model it has to grow above 20% its initial size. However, the final size of the tumour under continuous therapy is limited by the effective carrying capacity of the resistant population, $k_{\text{Eff}}$ - the resistant population can not expand above this value. A simple steady state analysis shows that the effective carrying capacity is given by:

$$k_{\text{Eff}} = (1 - \hat{d}_R / \hat{r}_R) \tag{4}$$

Thus, if $k_{\text{Eff}} < 1.2 n_0$ a tumour can not progress under continuous therapy nor adaptive therapy (for an example see Figures S3 A & B). This is a consequence of the simplifying assumption that the environmental carrying capacity, $K$, remains constant over time. In reality, the carrying capacity will expand over time due to angiogenesis and so the tumour would still eventually progress. As a result, the model behaviour in this parameter domain is not representative of the underlying biology, and we did not consider these cases further. For discussion of a model which accounts for a dynamic carrying capacity due to angiogenesis during adaptive therapy, see [8] and [9].

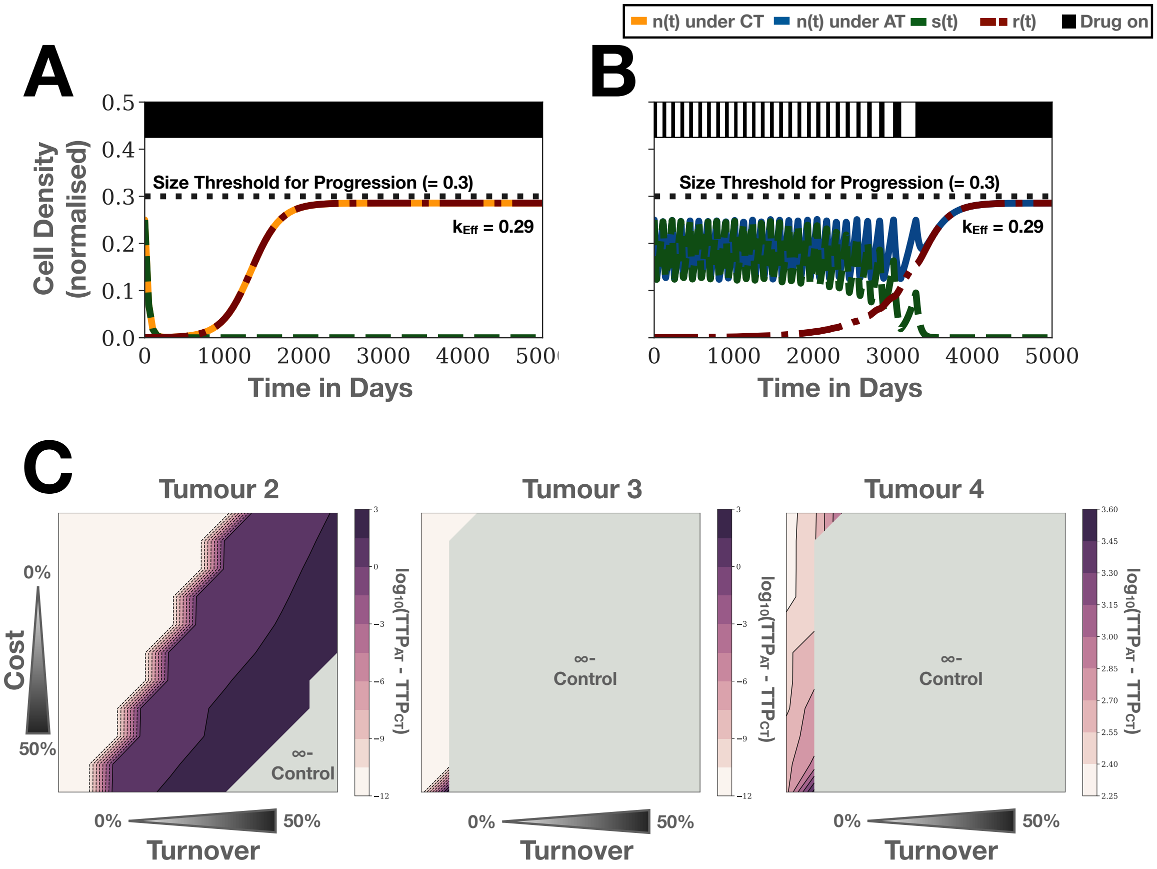


Figure S3: *As a consequence of the assumption that carrying capacity is constant the model allows for indefinite tumour control when* $(1 - \hat{d}_R/\hat{r}_R) < 1.2n_0$*. A: Example simulation illustrating indefinite control under continuous therapy for Tumour 1. While the tumour becomes completely resistant it can not grow above the 20% size increase required for progression, as the effective carrying capacity of the resistant population is too low. Parameters:* $n_0 = 25\%, f_R = 0.1\%, \hat{r}_R = 0.7, \hat{d}_R = 0.5$*. B: Same tumour as in A, but treated with adaptive therapy. Also in this case the tumour does not progress. C: The gain by adaptive therapy as a function of turnover and cost for Tumours 2-4. As Tumour 2 is far from carrying capacity* $(n_0 = 0.25)$ *and has a high resistance portion, strong turnover and/or big resistance costs are required to see significant benefits of adaptive therapy. Conversely, since they are close to K* $(n_0 = 0.75)$ *Tumours 3 & 4 are indefinitely controllable for a wide parameter regime.*

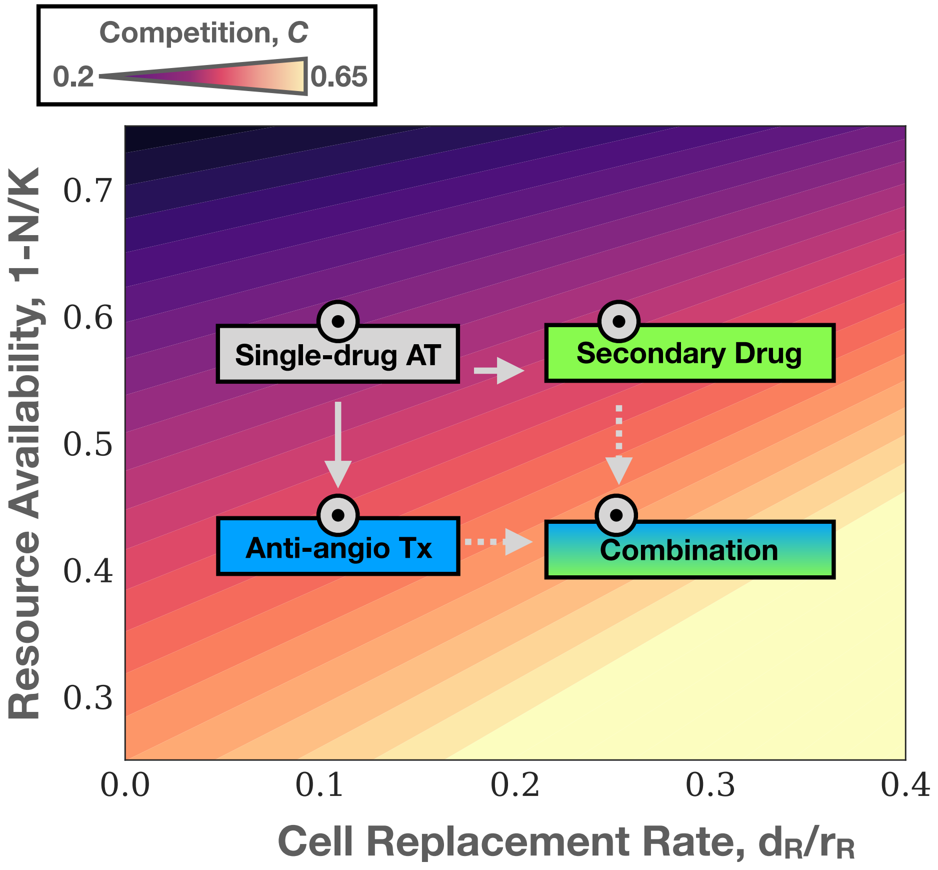


Figure S4: *Improving tumour control by increasing competition with a second drug which either reduces K or increases* $d_T/r_R$. *Here we show the intensity of competition at the start of treatment* ($C((1-f_R)n_0, f_R n_0, 1, \hat{d}_T, 1)$) *for the four cases considered in Figure 5E. Single drug adaptive therapy:* $n_0 = 0.4, f_R = 0.01, \hat{d}_T = 0.1$*; Anti-angiogenic Tx changes* $n_0$ *to 0.55; Secondary drug increases* $\hat{d}_T$ *to 0.25*

#### S6 Extension to other types of cost

Throughout the main body of our manuscript we assume that the resistance cost manifests itself in the proliferation rate of the cells. However, it is plausible that it might manifests itself in the carrying capacity or the death rate (see, for example, Figure S1). To study the dynamics of such manifestations of a cost we will extend our model to the following more general form:

$$\frac{dS}{dt} = r_S \left(1 - \frac{S+R}{K_S}\right)(1 - d_D D(t))\, S - d_S S, \tag{5}$$

$$\frac{dR}{dt} = r_R \left(1 - \frac{R+S}{K_R}\right) R - d_R R, \tag{6}$$

$$N(t) = S(t) + R(t), \tag{7}$$

In the main manuscript we show that the effective carrying capacity of the resistant population is a useful framework for understanding the impact of a cost in the growth rate. Thus, we will initially compare how $d_R$ and $K_R$ affect this equilibrium. Subsequently, we will show a set of simulations which demonstrates that when comparing different types of costs, we also need to take into account how each parameter affects the (density-independent) growth rate.

##### S6.1 Impact on the effective carrying capacity

The effective carrying capacity of the resistant population in this more general model is given by:

$$K_{\text{Eff}} = \left(1 - \frac{d_R}{r_R}\right) K_R \tag{8}$$

Equation (8) shows that costs in $d_R$ and $K_R$ also decrease the resistant population's effective carrying capacity, $K_{\text{Eff}}$, which will increase the benefit of AT, similar to what we have discussed for $r_R$. To compare the relative impact caused by each of the three types of cost, we define $R^*_r = \left(1 - \frac{d_R}{(1-c)r_R}\right) K_R$, $R^*_d = \left(1 - \frac{(1+c)d_R}{r_R}\right) K_R$, and $R^*_K = \left(1 - \frac{d_R}{r_R}\right)(1-c)K_R$ to be the population equilibria under a relative cost of $c$ ($> 0$) in $r_R$, $d_R$, and $K_R$, respectively. In Figure S5A-C we plot the three functions for three different values of the cell replacement rate $d_R/r_R$. When $d_R/r_R = 0$, only a cost in $K_R$ has an impact on the effective carrying capacity (Figure S5A). When we increase $d_R/r_R$ to 30%, small costs will have the greatest impact if they are in $K_R$, but if the costs are larger, they will have more impact in $r_R$ (Figure S5B). The transition occurs when the $R^*_r(c)$ and $R^*_K(c)$ lines intersect at $\frac{d_R}{r_R} = \frac{1-c}{2-c}$ (Figure S5B). As we will show below, this behaviour holds true for $0 < \frac{d_R}{r_R} < 0.5$. Finally, when $\frac{d_R}{r_R} > 0.5$, a cost in $r_R$ will have the greatest impact, followed by costs in $d_R$, and $K_R$, respectively.

We formally prove these observations in the following theorem:

**Theorem S6.1.** *Let* $R^*_r = \left(1 - \frac{d_R}{(1-c)r_R}\right) K_R$, $R^*_d = \left(1 - \frac{(1+c)d_R}{r_R}\right) K_R$, *and* $R^*_K = \left(1 - \frac{d_R}{r_R}\right)(1-c)K_R$ *denote the population equilibria under a relative cost of* $c$ ($> 0$) *in* $r_R$, $d_R$, *and* $K_R$, *respectively. Then:*

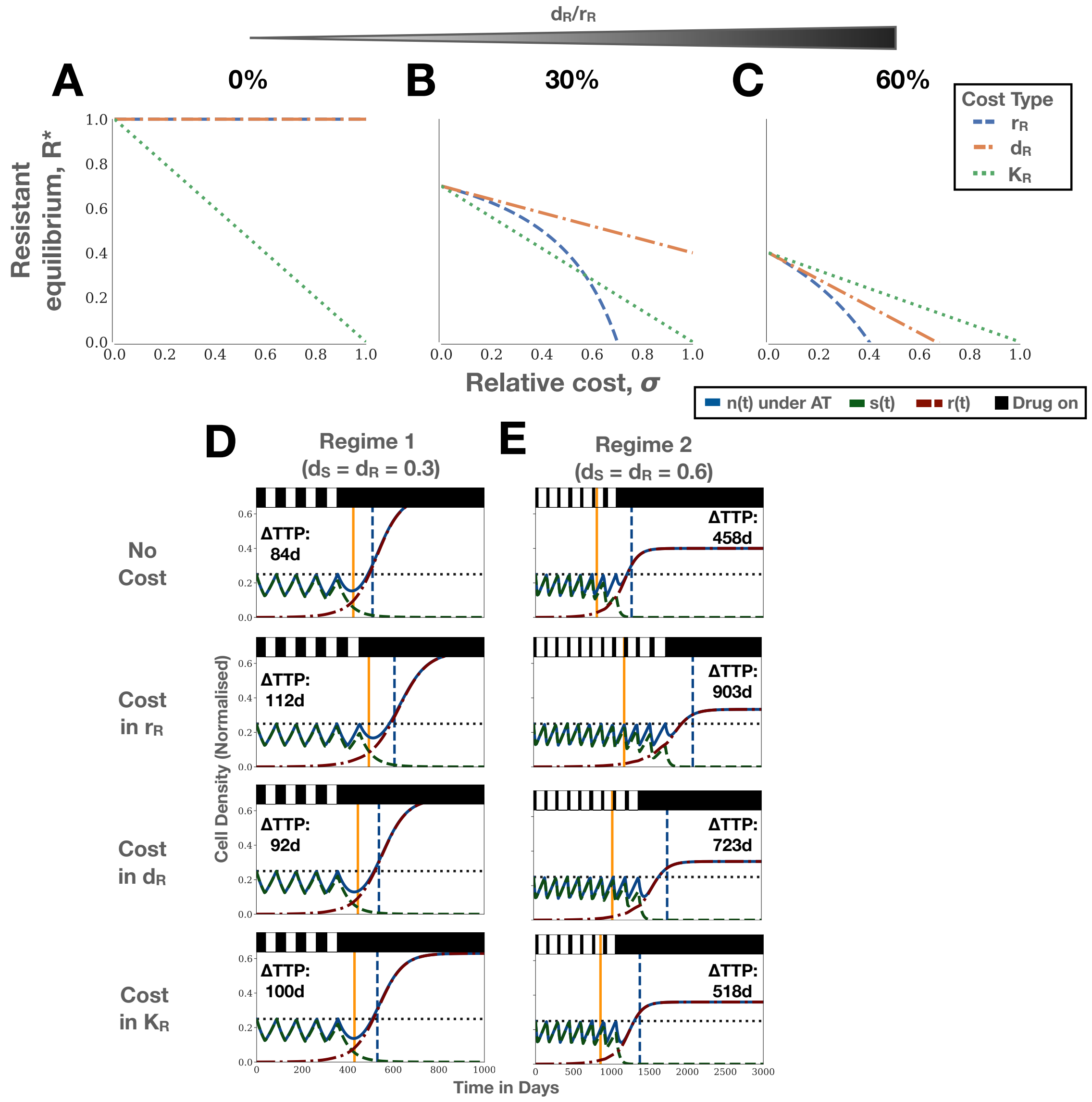


Figure S5: *Comparison of the impact of different manifestations of resistance costs. A-C) Plots of the resistant population equilibrium under costs in $r_R$ ($R_r^*(c)$), $d_R$ ($R_d^*(c)$), and $K_R$ ($R_K^*(c)$), respectively. The smaller the value of $R^*$, the longer the achievable tumour control. Note that when $R^* \leq 0$ the resistant population will always go extinct. A) When there is no turnover, only a cost in $K_R$ has an impact on $R^*$. B) For values of $d_R/r_R < 0.5$, there are two possible cases. For $c \leq 1 - \frac{1}{r_R/d_R - 1}$, a cost in $K_R$ has the greatest impact. Otherwise a cost in $r_R$ has a greater impact. C) For $d_R/r_R > 0.5$, a cost in $r_R$ always has the greatest impact, followed by costs in $d_R$ and $K_R$. D & E ) Treatment dynamics under AT for Tumour 1 from Figure 3 ($(n_0, f_R) = (25\%, 0.1\%)$) without a cost and with a 10% cost in each of $r_R$, $d_R$ and $K_R$, for two values of the replacement rate. D) Simulations corresponding to the parameters in B, in which a cost in $K_R$ has the greatest effect on $K_{Eff}$. Interestingly, we see that here, due to the small value of $n_0$, a cost in $r_R$ has the biggest impact on TTP. E) Simulations corresponding to C. Here, the results match those expected from Theorem S6.1, with again a cost in $r_R$ being the most impactful. This illustrates that the comparison of different types of costs also depends on the density of sensitive cells in the tumour.*

1. *The impact of a cost in $r_R$ will always be greater than the impact of a cost in $d_R$. That is, $R_r^* > R_d^*, \forall c \in (0,1)$.*

2. *Provided $d_R < \frac{c}{1+c} r_R$, the impact of a cost in $K$ will be greater than the impact of a cost in $d_R$ ($R_K^* > R_r^*$). Otherwise a cost in $d_R$ will have a greater impact.*

3. *The impact of a cost in $K$ will be greater than the impact of a cost in $r_R$ ($R_K^* > R_r^*$) provided $d_R < \frac{1-c}{2-c} r_R$. Otherwise a cost in $r_R$ will have a greater impact.*

*Proof.* Claim 1: Comparing $R_r^*$ and $R_d^*$ we find:

$$\begin{aligned}
&R_r^* > R_d^* \\
\Leftrightarrow &\left(1 - \frac{d_R}{(1-c)r_R}\right) K_R > \left(1 - \frac{(1+c)d_R}{r_R}\right) K_R \\
\Leftrightarrow &(1+c) < \frac{1}{1-c} \\
\Leftrightarrow &c^2 > 0,
\end{aligned}$$

so that $R_r^* > R_d^*, \forall c \in (0,1)$.

Claim 2: Similarly, comparing $R_K^*$ and $R_d^*$ gives:

$$\begin{aligned}
&R_K^* > R_d^* \\
\Leftrightarrow &\left(1 - \frac{d_R}{r_R}\right) (1-c)K_R > \left(1 - \frac{(1+c)d_R}{r_R}\right) K_R \\
\Leftrightarrow &- c(1+c)d_R - r_R(1+c) + c\,r_R(1+c) > -r_R \\
\Leftrightarrow &d_R < \frac{c}{1+c} r_R,
\end{aligned}$$

where we have skipped some steps for brevity.

Claim 3: Finally, comparing $R_K^*$ and $R_r^*$ yields:

$$\begin{aligned}
&R_K^* > R_r^* \\
\Leftrightarrow &\left(1 - \frac{d_R}{r_R}\right) (1-c)K_R > \left(1 - \frac{d_R}{(1-c)r_R}\right) K_R \\
\Leftrightarrow &- (1-c^2)\frac{d_R}{r_R} - c(1-c) > -\frac{d_R}{r_R} \\
\Leftrightarrow &d_R < \frac{1-c}{2-c} r_R,
\end{aligned}$$

where we again have skipped some steps for brevity. □

There are two important conclusions to draw from this analysis. Firstly, a cost in the growth rate will always have a greater impact than the same relative increase in the death rate (unless $d_R = 0$). Secondly, the relative importance of each of the three parameters depends on the ratio $d_R/r_R$.

##### S6.2 Comparison of the AT treatment dynamics under different types of cost

How do the different possible manifestation of a cost affect treatment dynamics? In Figures S5C & D, we show the AT treatment dynamics for a 10% cost in each of $r_R$, $d_R$ and $K_R$ for Tumour 1. Firstly, we consider the regime $d_R/r_R < 0.5$ in which a cost in $K_R$ will have the greatest impact on $K_{\text{Eff}}$. Interestingly, we find that - contrary to our expectations from Theorem S6.1 - a cost in $r_R$ has the largest impact on TTP (Figure S5D). As we can see from Figure S5D, the resistant population takes longer to expand and more AT cycles can be completed when there is a cost in $r_R$ compared to when there is a cost in $K_R$ (4 vs 5 cycles). To explain this, we rewrite Equation (6) as:

$$\frac{1}{r_R}\frac{dR}{Rdt} = \underbrace{\left(1 - \frac{d_R}{r_R}\right)}_{\substack{\text{Density-Independent}\\\text{Expansion}}} - \underbrace{\frac{S+R}{K_R}}_{\substack{\text{Density-Dependent}\\\text{Inhibition}}} . \tag{9}$$

We see that the benefit from the reduced carrying capacity depends on the density of sensitive cells. Is the tumour far from carrying capacity, as is the case here ($S \leq 0.25$), the benefit is small as the competition isn't very strong. In contrast, costs in $r_R$ (and $d_R$) have an impact *independent* of the proximity to carrying capacity.

In the second case, we assume that the replacement rate is greater than 0.5. While it is probably not biologically realistic, we include this case for completeness. We find that here the order in which cost extends TTP coincides with what would be expected from Theorem S6.1 (Figure S5E). This is because the costs in $r_R$ and $d_R$ reduce both the density dependent, and the independent-growth rate more than the cost in $K_R$ (Figure S5E).

This analysis demonstrates again that in order to understand the impact of a resistance cost we have to consider not only the cost itself, but also the context in which it occurs (here replacement rate and initial proximity to carrying capacity). Furthermore it shows that the two ways we discuss for improving AT (decreasing $K_R$ or increasing $d_R/r_R$) will amplify resistance costs regardless of their type.

[3] G. G. Steel, K. Adams, and J. C. Barrett, "Analysis of the cell population kinetics of transplanted

tumours of widely-differing growth rate," *British Journal of Cancer*, vol. 20, no. 4, pp. 784–800, 1966.

[4] G. G. Steel, "Cell loss as a factor in the growth rate of human tumours," *European Journal of Cancer (1965)*, vol. 3, no. 4-5, pp. 381–387, 1967.

[5] G. G. Steel, "CELL LOSS FROM EXPERIMENTAL TUMOURS," *Cell Proliferation*, vol. 1, no. 3, pp. 193–207, 1968.

[6] D. A. Rew and G. D. Wilson, "Cell production rates in human tissues and tumours and their significance. Part 1: An introduction to the techniques of measurement and their limitations," 2000.

[7] D. A. Rew and G. D. Wilson, "Cell production rates in human tissues and tumours and their significance. Part II: clinical data," 2000.

[8] R. A. Gatenby, A. S. Silva, R. J. Gillies, and B. R. Frieden, "Adaptive therapy," *Cancer Research*, vol. 69, pp. 4894–4903, jul 2009.

[9] K. Bacevic, R. Noble, A. Soffar, O. W. Ammar, B. Boszonyik, S. Prieto, C. Vincent, M. E. Hochberg, L. Krasinska, and D. Fisher, "Spatial competition constrains resistance to targeted cancer therapy," *Nature communications*, vol. 8, no. 1, pp. 1–15, 2017.
